# The “good, the bad and the double-sword” effects of exposure to MPs and their organic additives on N_2_-fixing bacteria

**DOI:** 10.1101/2020.07.21.210740

**Authors:** Víctor Fernández-Juárez, Xabier López-Alforja, Aida Frank-Comas, Pedro Echeveste, Antoni Bennasar-Figueras, Guillem Ramis-Munar, Rosa María Gomila, Nona S. R. Agawin

**Affiliations:** Marine Ecology and Systematics, Department of Biology, University of the Balearic Islands, Palma de Mallorca, Spain.; Instituto de Ciencias Naturales Alexander von Humboldt, Universidad de Antofagasta, Antofagasta, Chile.; Grup de Recerca en Microbiologia, Departament de Biologia, Universitat de les Illes Balears, Palma de Mallorca, Spain.; Celomic unit of the University Institute of Research in Health Sciences of the Balearic Islands, Palma de Mallorca, Spain.; Servicio Científico-Técnicos, University of the Balearic Islands, Palma de Mallorca, Spain.

**Keywords:** microplastics, organic additives, diazotrophs, *Posidonia oceanica* and N_2_-fixation.

## Abstract

The accumulation of microplastics (MPs) pollution at depths suggests the susceptibility of benthic organisms (*e.g.* seagrasses and their associated macro- and micro-organisms) to the effects of these pollutants. Little is known about the direct effects of MPs and their organic additives on marine bacteria, *e.g.* in one of the most ecologically significant groups, the diazotrophs or N_2_-fixing bacteria. To fill this gap of knowledge, we exposed marine diazotrophs found in association with the endemic Mediterranean seagrass *Posidonia oceanica* to pure MPs which differ in physical properties (*e.g.* density, hydrophobicity and/or size), namely, polyethylene (PE), polypropylene (PP), polyvinyl chloride (PVC) and polystyrene (PS) and to their most abundant associated organic additives (*e.g.* fluoranthene, 1,2,5,6,9,10-hexabromocyclododecane [HBCD] and dioctyl-phthalate [DEHP]). Growth, protein overexpression, direct physical interactions between MPs and bacteria, phosphorus (P) acquisition mechanisms and N_2_-fixation rates were evaluated. Our results show species-specific responses of the autotrophic and heterotrophic N_2_-fixing bacteria tested and the responses were dependent on the type and concentration of MPs and additives. N_2_-fixing cyanobacteria were positively affected by environmental and high concentrations of MPs (*e.g.* PVC), as opposed to heterotrophic strains, that were only positively affected with high concentrations of ∼120 µm-size MPs (detecting the overexpression of proteins related to plastic degradation and C-transport), and negatively affected by 1 µm-size PS beads. Generally, the organic additives (*e.g.* fluoranthene) had a deleterious effect in both autotrophic and heterotrophic N_2_-fixing bacteria and the magnitude of the effect is suggested to be dependent on bacterial size. We did not find evidences that specific N_2_-fixation rates were significantly affected by exposure to MPs, albeit changes in bacterial abundance can affect the bulk N_2_-fixation rates. In summary, we reported for the first time, the beneficial (the “good”), deleterious (the “bad”) and/or both (the “double-sword”) effects of exposure to MPs and their organic additives on diazotrophs found in association with seagrasses.

## **1.** Introduction

Marine bacteria are major players of biogeochemical cycles and food webs in the oceans (Anderson, 2018; Renn, 1937). One of the most ecologically significant groups of marine bacteria, the diazotrophs or N_2_-fixing bacteria, can convert atmospheric dinitrogen gas (N_2_) to readily usable form of dissolved inorganic nitrogen, *i.e.* ammonia (NH_3_). The biological N_2_-fixation (BNF) carried out by these microorganisms is the largest source of new nitrogen (N) in the oceans compensating the losses of N through denitrification (Browning et al., 2017). BNF is also essential in benthic systems (Sohm et al., 2011), such as in *Posidonia oceanica* seagrass beds (Agawin et al., 2017). This marine phanerogam is an endemic plant in the Mediterranean Sea with important ecological functions (*e.g.* maintains clear and oxygenated waters, supports high diversity of macro- and micro-organisms, with high primary production, can act as carbon sink and is a hot-spot of N_2_-fixation processes) (Campagne et al., 2015). The N_2_-fixing communities associated with *P. oceanica* may potentially supply the entire N demand for these plants (Agawin et al., 2019, 2017, 2016), and thus the N_2_-fixers are important in maintaining their high productivity and consequently the numerous ecological services that these plants offer (Agawin et al., 2019). However, seagrasses are subject to different anthropogenic threats (*e.g.* eutrophication, climate change and anchoring of boats) but the effect of marine plastic pollution on these communities remains to be investigated.

Marine coastal ecosystems are the most impacted zones by the pollution of plastics. Up to 10 million tons of plastic enters annually in the oceans (Almroth, 2019). This oceanic “soup” of plastic is composed of different particle size: macroplastics (> 250 mm), mesoplastics (1-25 mm), microplastics (MPs) (1-1000 μm) and nanoplastics (NPs) (< 1 μm) (Hartmann et al., 2019). Considering that approximately 10% of the world’s coastal population are living in the coastal areas of the Mediterranean Sea, it is one of the most polluted seas by plastics (de Haan et al., 2019). Available data from 2010 to 2019 revealed that an average of 1.94 x10^6^ ± 5.64 x10^5^ MPs Km^-2^ float on the surface of the Mediterranean Sea. The most impacted area is the Adriatic Sea, with 1.07 x10^7^ ± 3.14 x10^6^ MPs Km^-2^, followed by the Balearic Sea with 3.11 x10^5^ ± 1.35 x10^5^ MPs Km^-2^ (supplementary Fig. 1).

The most abundant polymers (at the macro- and micro-size scale) on the surface of the oceans and the Mediterranean Sea are polyethylene (PE) followed by polypropylene (PP) and then by others such as polyvinyl chloride (PVC) or polystyrene (PS) (Suaria et al., 2016). The PE and PP represent up to 68% of the total MPs waste in the Mediterranean Sea (Suaria et al., 2016). The abundance of plastic particles declines exponentially with depth according to their densities, resulting in low density polymers (*e.g.* PE and PP) predominating in the surface waters and higher density polymers (*e.g.* PS) in the deeper areas (Erni-Cassola et al., 2019). However, evidences suggest that much of the small plastic particles at the surface, independently of their density, end up in sediments by transport mechanisms (Reisser et al., 2015; Urbanek et al., 2018). MPs have associated chemical additives (usually organic in nature) that have been added to them to improve their chemical properties and these low molecular weight additives can leach from the plastic polymers and can also be sorbed onto them (Bakir et al., 2014). Therefore, MPs can be sources and vectors for organic pollutants (*e.g.* plastics additives) which are deleterious for marine organisms (Hahladakis et al., 2018). Thus, the accumulation of MPs pollution at depths suggests the susceptibility of benthic organisms (such as *P. oceanica* and their associated organisms) to the effects of these pollutants.

In eukaryotic microorganisms, the deleterious effects of plastics have already been described (Cole and Galloway, 2015; Nelms et al., 2018; Wang and Zheng, 2008), but studies investigating the direct effects of plastics on marine prokaryotic organisms are scarce (Bryant et al., 2016; Harrison et al., 2011; Romera-Castillo et al., 2018; Tetu et al., 2019), and nothing is known on the direct effect of MPs and their plastic additives on the growth, biochemistry, nutrient acquisition mechanisms and N_2_-fixation rates of marine diazotrophs. The few studies that have been done on other microorganisms are focused on plastic degradation and biofilm formation (Urbanek et al., 2018). Other studies have reported changes of the microbial communities associated with the floating plastics through metagenomic analysis (Yang et al., 2019), suggesting that the response to plastic pollution can be species-specific. Tetu et al. (2019) also revealed the importance of concentration levels of leached plastic in the most abundant photosynthetic bacteria in the oceans, *Prochlorococcus*. Considering these previous results, experimental studies investigating the effect of MPs and their additives should take into account the response of different test species and different concentration levels of the pollutants. Moreover, MPs have varying physical properties (*e.g.* density, hydrophobic or size) which may affect the response of microorganisms.

Here, we report for the first time the responses of different species of marine diazotrophs found in association with *P. oceanica* to different concentrations of MPs (*i.e.* PE, PP, PVC and PS) and their most predominant organic chemical additives (fluoranthene, 1,2,5,6,9,10-hexabromocyclododecane [HBCD] and dioctyl-phthalate [DEHP]). Our results show beneficial, detrimental or both effects, depending on the species tested and the type and concentrations of MPs and additives added.

## 2. Materials and Methods

### 2.1 Culture strains tested

Five N_2_-fixing species (two cyanobacteria and three heterotrophic bacteria) found in association with *P. oceanica* were selected as models to describe the effect of MPs/organic additives in diazotrophs. Four of them were previously detected through *nifH* analysis (*Halothece* sp., *Fischerella muscicola*, *Marinobacterium litorale* and *Pseudomonas azotifigens*) (Agawin et al., 2017). We selected the most related culturable strains: *Halothece* sp. PCC 7418, *F. muscicola* PCC 73103, *M. litorale* DSM 23545 and *P. azotifigens* DSM 17556^T^. The first two autotrophic strains, the unicellular *Halothece* sp. and the filamentous heterocyst-forming *F. muscicola* were obtained from the Pasteur culture collection of cyanobacteria (PCC). The heterotrophic bacteria (γ-proteobacteria) *M. litorale* and *P. azotifigens* were obtained from DSMZ-Deutsche sammlung von mikroorganismen und zellkulturen GmbH (DSMZ). In addition, we isolated one diazotrophic heterotrophic γ-proteobacteria from the roots of *P. oceanica*, identified as *Cobetia* sp. This strain was isolated by triturating the roots of *P. oceanica* in Tris-EDTA 1 mM pH 7.5 and cultured on N-free media and tested positive for *nifH* gene by polymerase chain reaction (PCR) and for acetylene reduction assay (ARA). By 16S analysis, the strain was identified as *Cobetia* sp. (UIB 001). Prior to the experiments, the cells were acclimatized and cultured in their respective optimal culture media to achieve their exponential phase. Culture media were composed of synthetic sea water medium (ASN-III) + Turks island salts 4X for *Halothece* sp., BG11^0^ for *F. muscicola* and marine broth (MB) for the rest of the heterotrophic bacteria (Stanier et al., 1979). The cells were acclimated at 25 °C at 120 r.p.m in a rotatory shaker with a photoperiod of 12-h (hours) dark:12-h light under low intensity fluorescent light (30 μE m^-2^ s^-1^).

### 2.2 Experimental culture conditions

All the experiments were performed in triplicate in a modified artificial sea water (ASW) medium (L^-1^: 25.0 g NaCl, 1.0 g MgCl_2_·6H_2_O, 5.0 g MgSO_4_·7H_2_O, 0.7 g KCl, 0.15 g CaCl_2_·2H_2_O, 0.1 g KBr, 0.04 g SrCl_2_·6H2O and 0.025 g H_3_BO_3_), 1 mL^-1^ trace metal [L^-1^: 2.86 g H_3_BO_3_, 1.81 g MnCl_2_·4H_2_O, 0.22 g ZnSO_4_·7H_2_O, 0.39 g NaMoO_4_·2H_2_O, 0.079 g CuSO_4_·5H_2_O and 0.049 g Co(NO_3_)_2_·6H2O], glucose (final concentration 0.1% [v/v]) at pH 7 and with the respective MPs and/or organic additives. Inorganic phosphorus (PO_4_^3-^, 0.04 g L^-1^ K_2_HPO_4_), iron (Fe, 0.006 g L^-1^ferric citrate) and inorganic nitrogen (NH_3_, 0.5 g L^-1^ NH_3_Cl) were added in function of the response variable measured as we described below. Bacteria at their exponential phase were inoculated in the treatments (supplementary Table 1) and incubated for 72 h under the same conditions as when they were previously acclimatized (*i.e.* at 25 °C, 120 r.p.m in a rotatory shaker with a photoperiod of 12 h dark:12 h light, under low intensity fluorescent light, 30 μE _m_^-2^ _s_^-1^_)._

The pure MPs (low-density polyethylene [PE] with size 109 ± 6.29 μm, polypropylene [PP] with size 90 ± 7.56 μm and low-density polyvinyl chloride [PVC] with size 164 ± 8.03 μm) and organic additives (fluoranthene, 1,2,5,6,9,10-hexabromocyclododecane [HBCD] and dioctyl-phthalate [DEHP]) used were obtained from Sigma-Aldrich. In addition, we used fluorescent polystyrene (PS)-based latex beads (Fluoresbrite® YG Microspheres 1.00 µm, Polysciences, Inc.) as the lowest sized MPs, based on the definition of MPs (Hartmann et al., 2019). Stock solution of MPs were made at 100 mg mL^-1^, resuspending the MPs (previously UV-sterilized for 15 min) in acetone (98% [v/v]) to avoid aggregation of MPs and for easier manipulation of the workings solutions. In order to prevent chemical damages of MPs by acetone, the stock solution was rapidly diluted to working solutions of 1 mg mL^-1^ in ASW. The organic additives, *i.e.* fluoranthene and HBCD, were initially prepared in 1 mg mL^-1^ in absolute ethanol and acetone (98% [v/v]), respectively. For the DEHP additive that was in liquid form, a stock solution of 1 mg mL^-1^ was also prepared. We diluted these stock solutions to working solutions of 3 mg L^-1^ in ASW. Fluorescent PS beads were sterilized following the manufacturer’s instructions, and the different concentrations in ASW were made from a stock solution of 4.55 x10^10^ particles mL^-1^. The controls were made with the respective amounts of acetone and/or ethanol (without any MPs nor organic additives). All the treatments have ≤ 1% of acetone or ethanol to avoid any biological effect in the cells, and co-solvents effect in water (Schwarzenbach et al., 2002).

Experiments were performed in two parts as described in the supplementary Table 1, following the recommendations of Paul-Pont et al. (2018). In the first part, we made an overall screening of the five culture strains in sterile 2 mL well microplates to study their growth response under marine environmentally relevant concentrations of MPs and additives, with optimal nutrient conditions of PO_4_^3-^, Fe and NH_3_. We used five concentration for MPs (0, 0.01, 0.1, 1 and 100 μg mL^-1^) and organic additives (0, 0.3, 3, 30 and 300 μg L^-1^) (supplementary Table 1). Additional treatments combining MPs and plastic additives (*e.g*. PE-fluoranthene, PP-HBCD and PVC-DEHP) were done by combining the lowest and the highest number of MPs and additives to test for possible interacting effects of MPs and their organic additives (supplementary Table 1). In the second part, we selected two strains, one autotrophic (*Halothece* sp.) which was easy to manipulate and one heterotrophic (*Cobetia* sp.) to compare their responses to marine environmental concentrations (MPs: 100 μg mL^-1^ and additives: 300 μg L^-1^) with very high concentrations (as “the worst case scenario”) of MPs and additives (MPs: 1000 μg mL^-1^ and additives: 3000 μg L^-1^). In addition, we selected a concentration of 4.55 x10^6^ and 4.55 x10^7^ particles mL^-1^ for PS beads. We cultured the test bacteria in 50 mL falcon tubes under N^2^-fixing conditions (limiting NH_3_ [~ 0.15 mM] and with optimal PO_4_^3-^ and Fe) for growth, protein overexpression, microscopical analysis, PO_4_^3-^-uptake and N_2_-fixation assays, or under PO_4_^3-^-limiting conditions (with optimal NH_3_ and Fe) for alkaline phosphatase activity (APA) (supplementary Table 1).

### 2.3 Flow cytometry

Aliquots of cultures from all the experiments were taken initially and after 72 h of incubation and counted in fresh (without freezing nor fixing) with a Becton Dickinson FACS-Verse cytometer (Beckton & Dickinson, Franklin Lakes, New Jersey, USA). Fluorescent beads, BD FACSuite™ CS&T research beads (Beckton & Dickinson and Company BD Biosciences, San Jose, USA), were used as internal standards to calibrate the instrument. Cells were separated by combinations of the scatter plots of the flow cytometer parameters: forward scatter (FSC, reflecting cell size), side scatter (SSC, reflecting internal complexity of the cells) and/or fluorescein isothiocyanate filter (FITC, reflecting fluorescence, 488 nm excitation, 530/30 nm emission). For treatments with fluorescent PS beads, adhesion to the PS were counted using the FSC and FITC cytometer signals. Adhered bacteria were those with intermediate intensity fluorescence signals between the bacteria and the beads. A total of 10000 cells (or cells recorded in 30 seconds) were counted in each sample, and the results were expressed as cells mL^-1.^ Changes in cell abundance were calculated as the changes in cell concentrations after 72 h (between the final and the initial day of the experiment).

### 2.4 Protein identification: MALDI-TOF assay and protein structure prediction

Crude cell extracts following the methods described in Ivleva and Golden (2007) were done with the cultures of *Halothece* sp. and *Cobetia* sp. after 72 h of incubation (at the highest concentration treatment of MPs/additives and the control). The cells were centrifuged at 6000 x g, 5 minutes, washed with phosphate buffer saline (PBS), lysed with acid-washed glass beads (150-212 μm, Sigma-Aldrich) and resuspended again with PBS. The suspension was spun briefly at 1000 x g and the resultant supernatants were collected as the crude cell extract. Protein concentrations were determined with Bradford protein assay (Thermofisher), following the manufacturer’s instructions. The protein extracts were ran into polyacrylamide gels, 4-20% (p/v) Amersham ECL Gel (GE Healthcare, Chicago, Illinois, EEUU), using the ECL Gel Box system (GE Healthcare, Chicago, Illinois, EEUU) following the manufacturer’s instructions.

The different bands, detected only in *Cobetia* sp. exposed to high concentrations of MPs (1000 ug mL^-1^ of PE, PP and PVC) which did not appear at the control, were excised from the gel with a clean scalpel and sent to MALDI-TOF analysis. Each gel slice was cut into small pieces and then transferred to a clean and sterile Eppendorf tube. Protein identification was performed following Jaén-Luchoro et al. (2017). Briefly, bands were washed with ammonium bicarbonate (NH_4_HCO_3_) and acetonitrile (CH_3_CN) and dried afterwards. The gel band was rehydrated with pre-chilled trypsin (20 μg mL^-1^ in NH_4_HCO_3_ 50 mM) and NH_4_HCO_3_ solution. An extraction solution of CH_3_CN, trifluoroacetic acid (C_2_HF_3_O_2_) and water were added, and the extract was deposited on a MALDI-TOF MS plate (polished-steel, Bruker-Daltonics) and dried at room temperature. The sample was then analyzed with an Autoflex III MALDI-TOF-TOF (BrukerDaltonics) spectrometer using the software Compassflex series v1.4 (flexControl v3.4, flexAnalysis v3.4 and BioTools 3.2). The spectra were calibrated using the peptide calibration standard (BrukerDaltonics). The obtained mass spectra were used for the protein identification and the in-house database was created with the predicted protein sequence of the annotated genome of *Cobetia* sp. The search process was performed with the algorithm MASCOT (MatrixSciences).

Fasta sequence of the alcohol dehydrogenase (ADH) (detected through MALDI-TOF) was sent to the I-Tasser server for protein 3D-structure prediction (Zhang, 2008), with their domains previously checked in Pfam 32.0 (Finn et al., 2016). The predicted structure was send to the FunFOLD2 server for the prediction of protein–ligand interactions (Roche et al., 2013). In addition, we detected the potential sites of ligand or “pockets” through MetaPocket 2.0 (Huang, 2009). Finally, we predicted the orientation and position of protein-ligand complex between polyethylene glycol (PEG) and ADH, docking these with Swissdock (Grosdidier et al., 2011). All the structures were visualized by Pymol (DeLano, 2002).

### 2.5 Microscopic observations

At the final time (after 72 h), the five strains tested were placed onto elisa plate (Biolinea S.L) for inverted microscopy visualization of the physical interactions between bacterial cells and the MPs (*i.e.* PE, PP and PVC) at 100x objective (Leica DM IRB). For *Halothece* sp. and *Cobetia* sp., their interaction with PS fluorescent beads were visualized by confocal microscopy (Leica TCS SPE, Leica Microsystems). These cells were placed onto a clean glass microscope slide for confocal microscopy and a 100x objective was used to visualize the cells. Images were processed using the software Leica application suite (Leica Microsystems). Autofluorescence for *Halothece* sp. was observed with an excitation of 532 nm and emission of 555-619 nm. For *Cobetia* sp., the cells were stained with Sybr green (Sigma-Aldrich) or propidium iodide (Sigma) to visualize the cells. Visualization of the 1 μm PS fluorescent beads was done following the manufacturer’s instructions, with an excitation of 441 nm and emission of 485 nm.

### 2.6 P-metabolism analysis

Alkaline phosphatase activity (APA) was evaluated through fluorometric assay following the hydrolysis of the substrate (S) 4-methylumbelliferyl phosphate (MUF-P, Sigma-Aldrich) to 4-methylumbelliferyl (MUF). An end point enzymatic assay was conducted at the final day of the experiment, following Fernández-Juárez et al. (2019). The culture media was PO_4_^3-^ limited (but with optimal NH_3_ and Fe) to promote APA. Saturation curves of velocity (V, fmol MUF cell^-1^ h^-1^) vs substrate (S, μm) were made for each experimental condition for each of the two selected strains (*Halothece* sp. and *Cobetia* sp.). We used different concentrations of MUF-P: 0 μm, 0.05 μm and 5 μm of MUF-P. After 1 h incubation in darkness at room temperature, APA was measured in a microtiter plate that contained buffer borated pH 10 (3:1 of sample:buffer). The MUF production (fmol MUF cell^-1^ h^-1^) was measured with a Cary Eclipse spectrofluorometer (FL0902M009, Agilent Technologies) at 359 nm (excitation) and 449 nm (emission), using a calibration standard curve with commercial MUF (Sigma-Aldrich). The maximum velocity (V^max^) at saturating substrate concentrations was obtained from each plot of V vs S, using Lineweaver-Burk plot.

For the determination of inorganic PO_4_^3-^ concentrations for *Halothece* sp. and *Cobetia* sp. experiments, 1 mL of the culture was centrifuged for 15 minutes at 16000 x g under 4 °C. The bacteria-free clear supernatant was collected and used for PO_4_^3-^ determinations following standard spectrophotometric methods (Hansen and Koroleff, 1999). The PO_4_^3-^ concentrations in the culture media were determined at initial and final time (after 72 h). Specific PO_4_^3-^-uptake rates (pmol PO_4_^3-^ cell^-1^ d^-1^) were calculated as described in Fernández-Juárez et al. (2019) and Ghaffar et al. (2017):

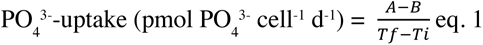

Where, *A* is the pmole PO_4_^3-^ cell^-1^ at the initial time (*Ti*) and *B* is the pmol PO_4_^3-^ cell^-1^ at the final time (*Tf*).

### 2.7 Determination of N_2_-fixation rates through acetylene reduction assay (ARA)

Rates of N_2_-fixation were measured for *Halothece* sp. A known volume of culture (8 mL) was sampled during the dark photoperiod and transferred to a closed hermetic vial and oxygen was flushed from sample through bubbling with N_2_ gas. A medium with saturated acetylene was injected at 20% (v/v) final concentration in each vial with a sterile syringe. Samples were incubated for 3 h at room temperature (24 °C) in the dark. After the 3 h incubation time, 10 mL of liquid were removed and transferred to Hungate tubes containing 1.25 mL of 20% trichloroacetic acid (Agawin et al., 2014). Prior to analysis with the gas chromatograph (GC), the Hungate tubes were incubated at 37 °C overnight in a water bath. Ethylene and acetylene gas from the gas phase of the Hungate tubes were determined using a GC (model GC-7890, Agilent Technologies) equipped with a flame ionization detector (FID). The column was a Varian wide-bore column (ref. CP7584) packed with CP-PoraPLOT U (27.5 m length, 0.53 mm inside diameter, 0.70 mm outside diameter, 20 μm film thickness). Helium was used as carrier gas at a flow rate of 30 mL min^-1^. Hydrogen and airflow rates were set at 30 mL min^-1^ and 365 mL min^-1^, respectively. The split flow was used so that the carrier gas flow through the column was 4 mL min^-1^ at a pressure of 5 psi. Oven, injection and detector temperatures were set at 52 °C, 120 °C and 170 °C, respectively. Ethylene produced was calculated using the equation in Stal (1988):

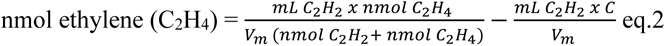

Where, *mL C_2_H_2_* is the amount of acetylene injected, *nmol C_2_H_2_* and *nmol C_2_H*^4^ are the amounts of acetylene and ethylene, respectively, as measured in a gas sample in the gas chromatograph. *Vm* the molar volume of an ideal gas and equals *RT/P* in which *R* is the molar gas constant, 8.31 Jmo1^-1^ K^-1^, *T* the temperature in degrees Kelvin, and *P* the pressure in N m^-2^ (pascals). At T = 273.15 K and P = 1 atm (101.32 N m^-2^), *Vm* equals 22.4 x10^-6^ mL nmol^-l^. At T = 293 K (20° and P = 1 atm), *Vm* equals 24.0 x10^-6^ mL nmol^-^ *C* is a correction factor to correct for the contamination of acetylene by ethylene and equals the molar ratio of C_2_H_4_ to C_2_H_2_. The acetylene reduction rates were converted to N_2_-fixation rates (nmol mL^-1^ h^-1^) using a factor of 4:1 (Jensen and Cox, 1983).

### 2.8 Statistical analysis

Normality of the data was checked using Shapiro-Wilk and Anderson-Darling for experiments with n < 30 and n > 30, respectively, and Levene’s test was used to assess the homogeneity of variance. Parametric analysis was used to analyze normally distributed data through 1- and 2-way ANOVAs with Bonferroni correction (Bonferroni post hoc test). For non-normally distributed data, Kruskal-Wallis rank sum non-parametric test as used. Dunn’s multiple comparison test with Bonferroni method was applied to determine the significant effects among different concentrations of MPs and additives. All analyses were done in R-Studio, R version 3.5.3 (2019-03-11).

## 3. Results

### 3.1 Effect of varying concentrations of MPs and additives in N_2_-fixers

#### 3.1.1 Concentrations under relevant concentrations in the marine environment

Polyethylene (PE) and PP within the range of 0-100 μg mL^-1^ did not significantly affect the growth of all cultures of diazotrophs tested (ANOVA or Kruskal-Wallis, p > 0.05, Figs. 1A-1B). PVC did not significantly affect the growth of the three heterotrophic diazotrophs (*Cobetia* sp., *Marinobacterium litorale* and *Pseudomonas azotifigens*) but for the two cyanobacterial diazotrophs (*Halothece* sp. and *Fischerella muscicola*), the PVC addition significantly enhanced their growth (ANOVA, p < 0.05), particularly at the highest PVC concentration compared to the control (Bonferroni, p < 0.05, Fig. 1C). The organic additives (*i.e.* fluoranthene and DEHP) significantly reduced the growth of *Halothece* sp. and *F. muscicola*, by 22-fold and 7-fold, respectively, at the highest concentrations compared to the control (Bonferroni, p < 0.05, Figs. 1D and 1F). On the contrary, the additive HBCD did not affect the growth of any species (ANOVA, p > 0.05, Fig. 1E), while the additive DEHP significantly enhanced the growth of the heterotrophic *P. azotifigens* by 4-fold at 30 μg L^-1^ compared to the control (Bonferroni, p < 0.05, Fig. 1F). PS beads at 4.55 x10^3^ particles mL^-1^ did not have any significant effect (data not shown).

**Figure 1.**
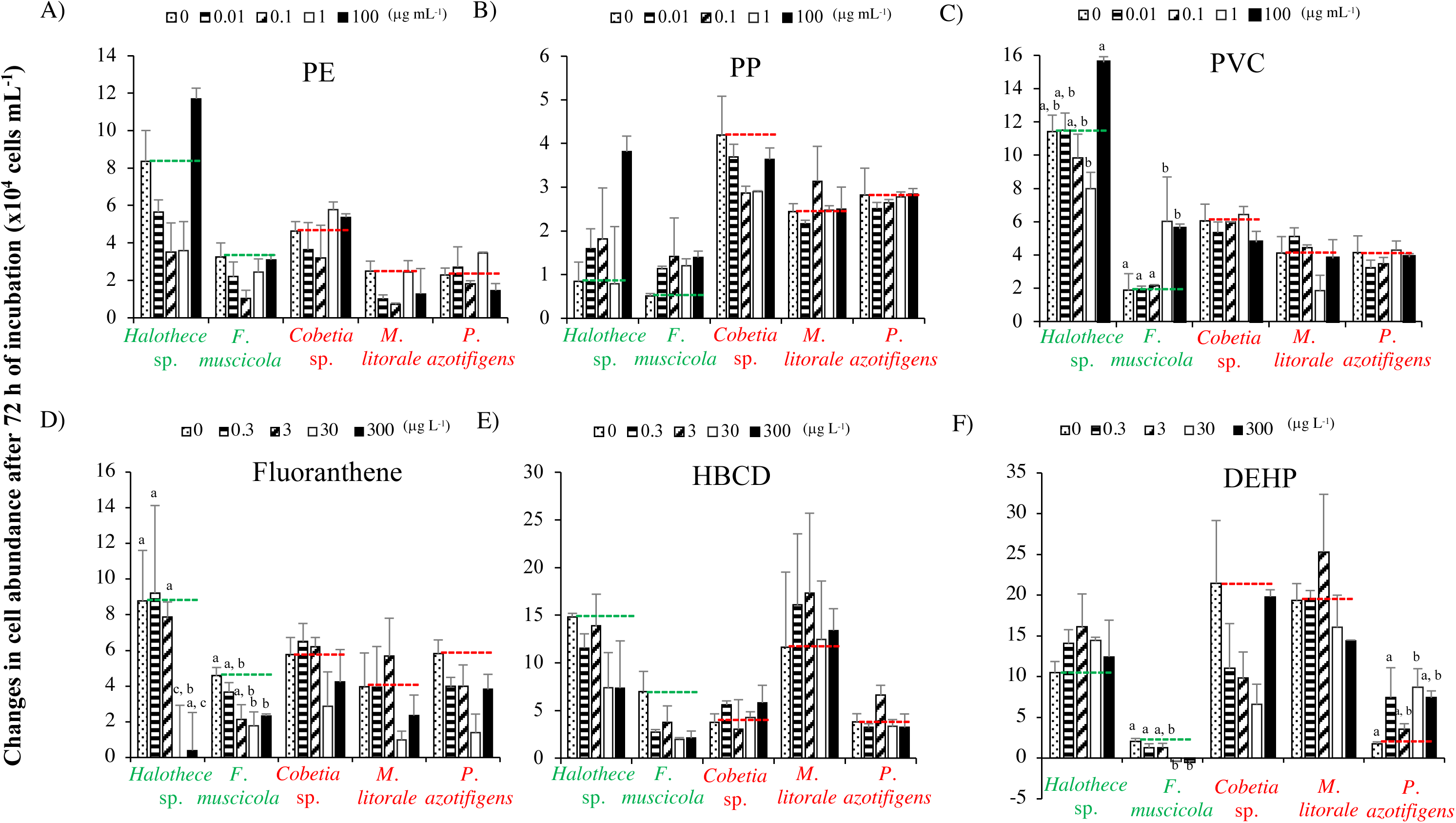
Growth responses of diazotrophs subject to MPs and organic additives under relevant concentrations in the marine environment. **A)** PE, **B)** PP, **C)** PVC, **D)** Fluoranthene, **E)** HBCD and **F)** DEHP. In **A-F)** values are the mean and the error bar is the standard error between the replicates. Letters indicate pairwise significant differences (p < 0.05) among the variables, only when significant differences were detected, using a posthoc test (Bonferroni or Dunn test) after ANOVA or Kruskal-Wallis over the whole dataset.

The interactions of each MP with their most common additive showed differing effects on the growth of diazotrophs (Fig. 2). The treatment PE + fluoranthene (at low and high concentrations) consistently decreased the growth of all diazotrophs tested (ANOVA or Kruskal-Wallis, p < 0.05, Fig. 2). The treatment PP + HBCD did not have a significant effect on the species tested (ANOVA or Kruskal-Wallis, p > 0.05, Fig. 2), while PVC + DEHP (at low and high concentrations) enhanced the growth of the heterotrophs *Cobetia* sp. and *P. azotifigens*, while it decreased the growth of *M. litorale* (Bonferroni test, p < 0.05, Fig. 2).

**Figure 2.**
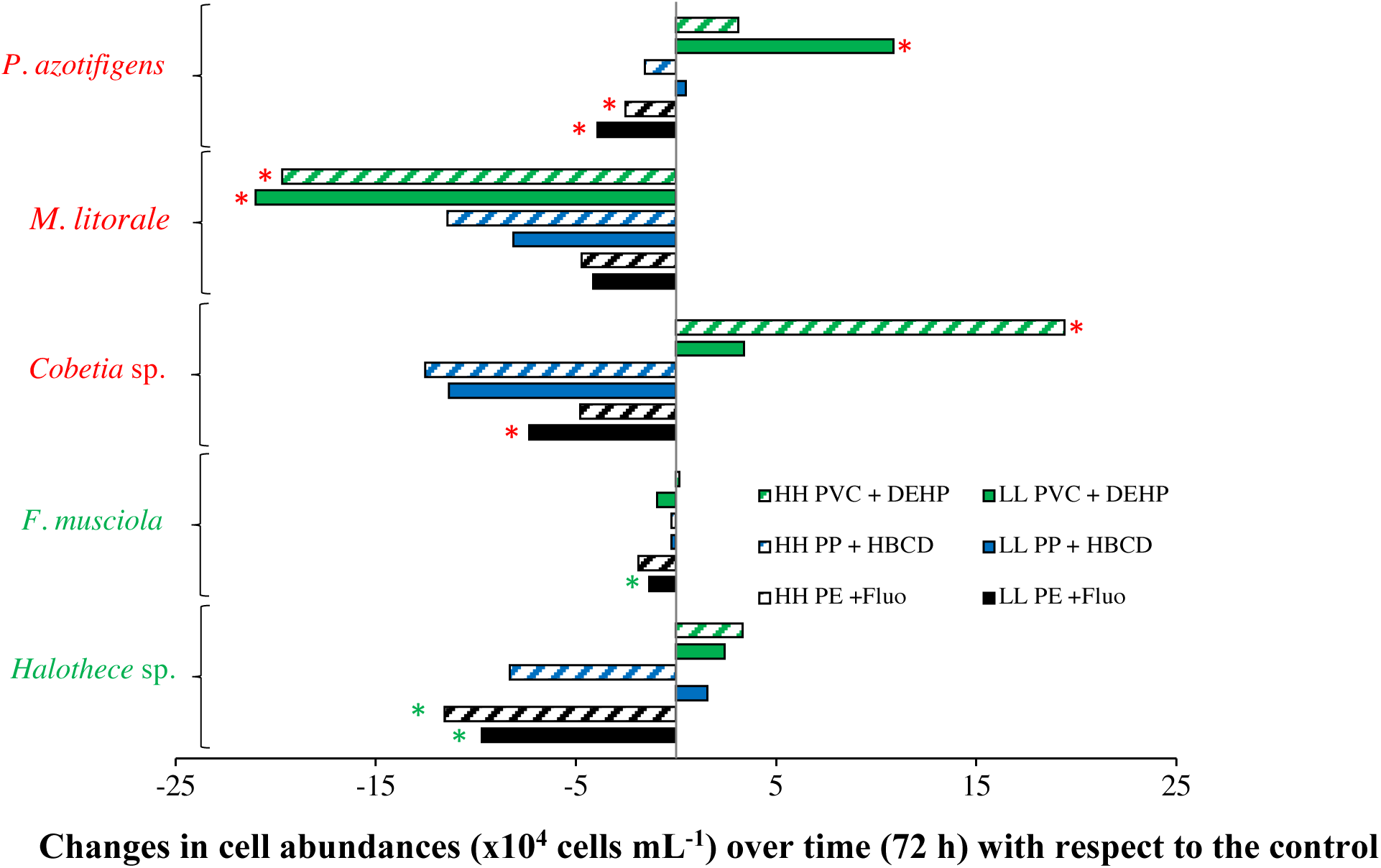
Growth responses with respect to control of the diazotrophs tested subject to MP-additive interactions (PE + fluoranthene, PP **+** HBCD and PVC + DEHP). “L” and “H” represent low and high concentrations of MPs (0.01 and 100 μg mL^-1^, respectively) and plastic additive (0.3 and 300 μg L^-1,^ respectively). Values are the mean and asterisks (*) indicate pairwise significant differences (p < 0.05) with the control, using a posthoc test (Bonferroni or Dunn test) after ANOVA or Kruskal-Wallis over the whole dataset.

#### 3.1.2 Response of MPs pollution under very high concentrations “worst case scenario”

##### 3.1.2.1 Effect of MPs (with an average size of 120 μm) and organic additives

Fig. 3A shows that MPs at high concentrations (up to 1000 μg mL^-1^) significantly affected the growth (ANOVA or Kruskal-Wallis, p < 0.05) of the two diazotrophic bacteria selected: the unicellular cyanobacteria *Halothece* sp. and the heterotroph *Cobetia* sp. High MPs concentrations increased bacterial population, especially with the addition of PVC in both strains compared with the control (Bonferroni test or Dunn test, p < 0.05, Fig. 3A). Contrary to the effects of MPs, the addition of different types of organic additives (fluoranthene, HBCD and/or DEHP up to 3000 μg L^-1^) affected negatively the growth of *Halothece* sp. (Dunn test, p < 0.05, Fig. 3B). For the heterotrophic bacteria, different responses were observed, with fluoranthene showing the highest toxicity (Dunn test, p < 0.05, Fig. 3B). The DEHP additive (at high concentrations) and the interaction of the three MPs with the three plastic additives at low concentrations enhanced the growth of *Cobetia* sp. (Dunn test, p < 0.05, Fig. 3B).

**Figure 3.**
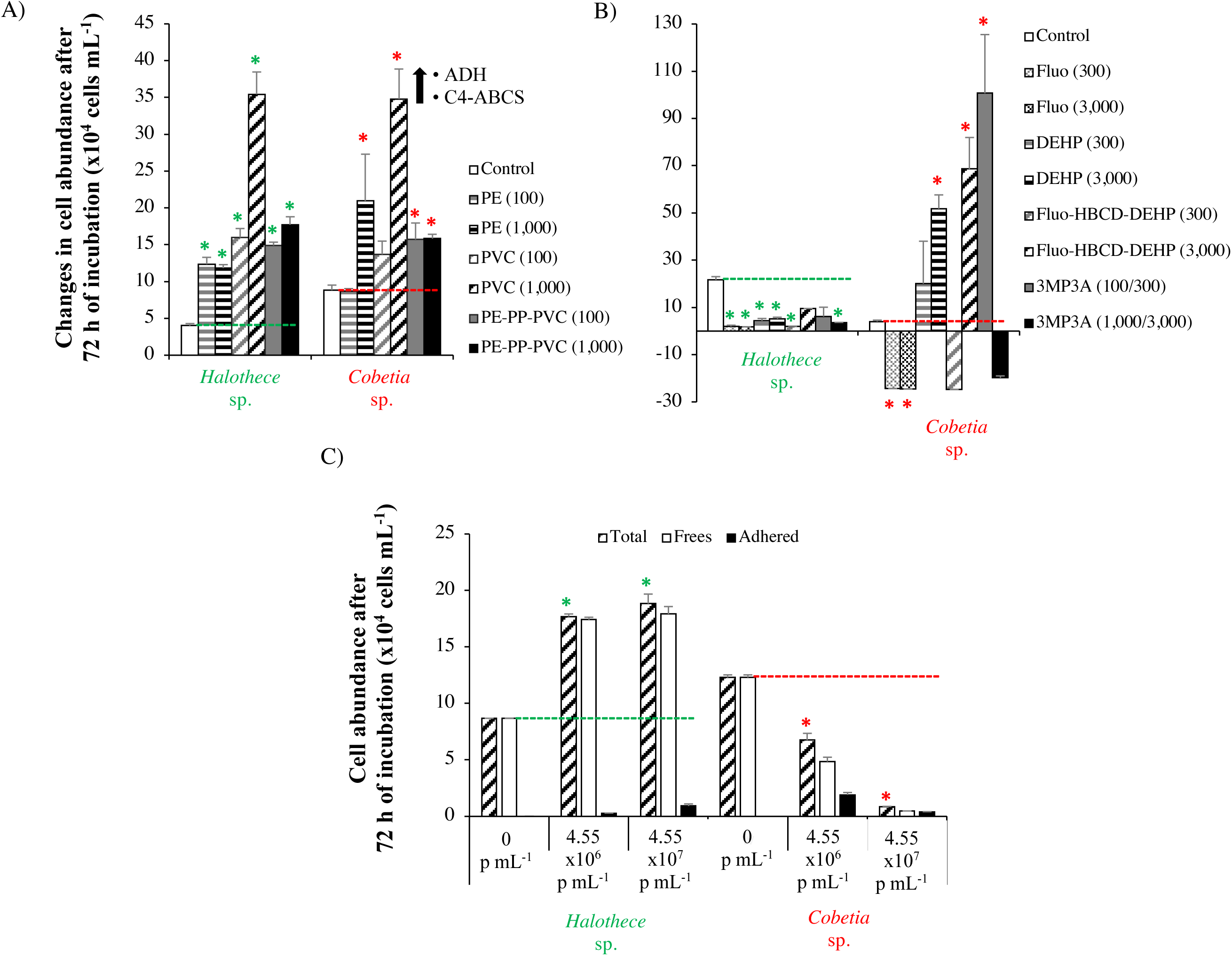
Growth responses after 72 h of incubation of diazotrophs subject to high concentrations of MPs and additives. **A)** Effect of MPs (PE, PP and/or PVC, 100/1000 µg mL^-1^). Black arrow indicates the protein overexpression detected through MALDI-TOF, only for *Cobetia* sp. The effect of PP alone with non-significant results is not included in the graph. **B)** Effect of organic additives (fluoranthene, HBCD and/or DEHP, 300/3000 µg L^-1^). The effect of HBCD alone with non-significant results is not included in the graph. **C)** Effect of PS beads. Total, free and adhered bacteria (to the beads) are differentiated. Values are the mean and the error bar is the standard error between the replicates. Asterisks (*) indicate pairwise significant differences (p < 0.05) with the control, using a posthoc test (Bonferroni or Dunn test) after ANOVA or Kruskal-Wallis over the whole dataset.

##### 3.1.2.2 Effect of smaller plastics (1 μm)

The effect of PS fluorescent beads (1 μm) was different between *Halothece* sp. and *Cobetia* sp. (ANOVA, p < 0.05, Fig. 3C). For *Halothece* sp., PS beads at 4.55 x10^6^ and 4.55 x10^7^ particles mL^-1^ enhanced significantly their growth (Bonferroni test, p < 0.05, Fig. 3C), with less than 5% of the cells being adhered to beads. On the contrary, PS beads significantly decreased the growth of *Cobetia* sp. (Bonferroni test, p < 0.05, Fig. 3C), with 40% to 87% of the cells being adhered to the PS beads.

##### 3.1.2.3 Changes in protein expression

Changes in protein expression were analyzed for *Halothece* sp. and *Cobetia* sp. when they were exposed to the high levels of MPs and additives. Overexpression of two proteins under the PE-PP-PVC condition (1000 μg mL^-1^) compared to the controls were detected only for *Cobetia* sp.: an alcohol dehydrogenase (ADH) of 342 aminoacids (aa) (in which is reported than can degrade polymers such as polyethylene glycol [PEG]) and a C4-dicarboxylate ABC transporter substrate-binding protein (C4-ABCS) of 329 aa (Fig. 3A). Our predicted ADH structure showed a nicotinamide-adenine-dinucleotide (NAD) dependence and a potential binding site with PEG (Figs. 4A-C). Protein overexpression were not detected after the addition of organic additives neither for *Halothece* sp. nor for *Cobetia* sp.

**Figure 4.**
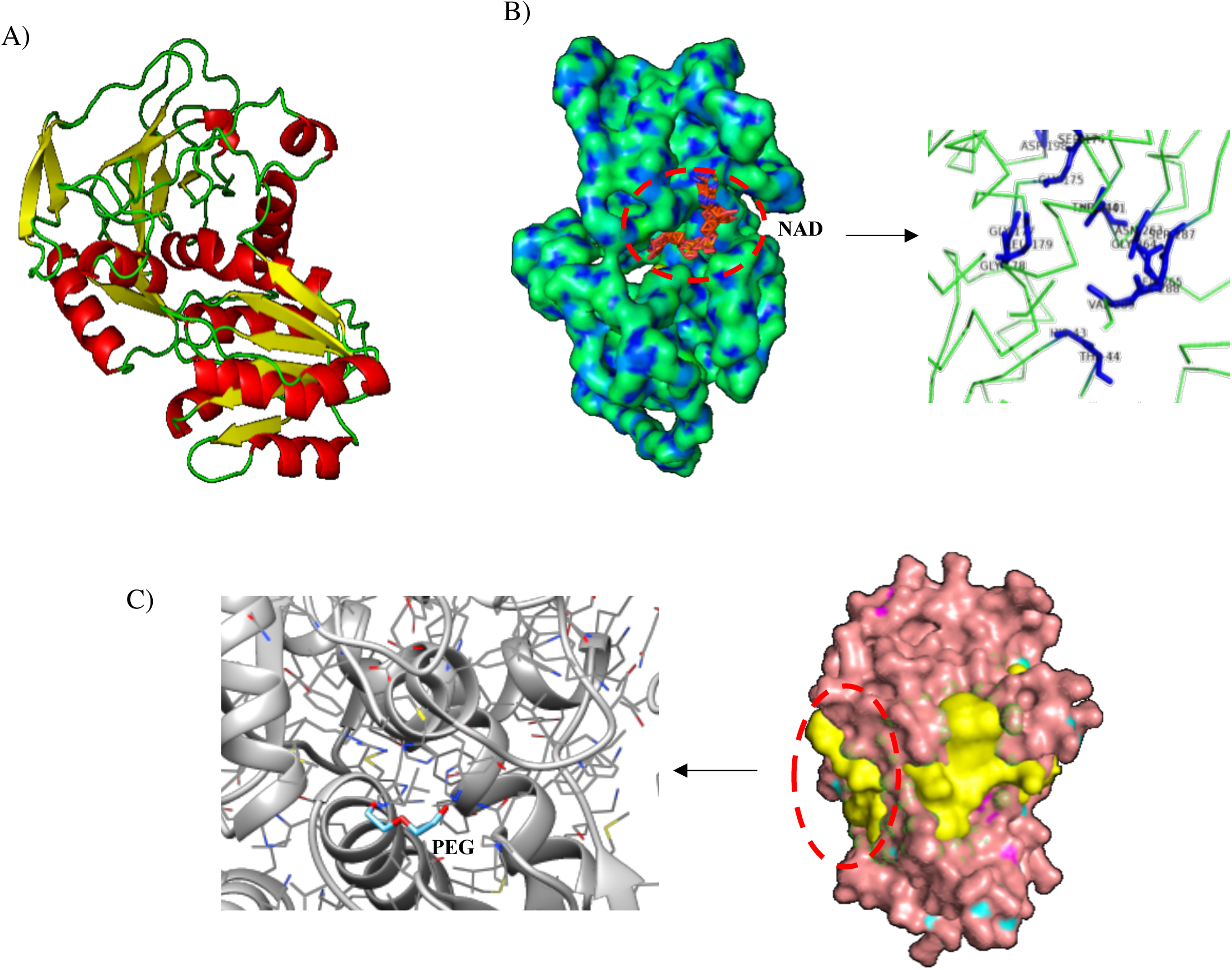
Structural analysis of alcohol dehydrogenase (ADH) from *Cobetia* sp. **A)** Protein prediction of ADH showing, **B)** nicotinamide-adenine-dinucleotide (NAD) dependence, displaying the ligand binding residues HIS 43, THR 44, SER 174, GLY 175, GLY 177, GLY 178, LEU 179, ASP 198, THR 240, ASP 241, ASN 263, GLY 264, LYS 265, SER 287, ILE 288 and VAL 289. **C)** The three of the most probable pockets of the ADH (potential sites of ligand binding, in yellow), and the docking site (prediction of the orientation and position of protein-ligand complex) between the polyethylene glycol (PEG) and ADH.

### 3.2 Physical interactions between MPs and diazotrophs

Cyanobacterial diazotrophs (*Halothece* sp. and *F. muscicola*) were not capable of adhering to the MPs, however, the bigger-size of *F. muscicola* cells (~ 7 μm per cell), which formed filaments, were trapped by MPs (Fig. 5A). Heterotrophic diazotrophs (*e.g. Cobetia* sp. or *M. litorale*) were adhered on MPs (Figs. 5B-E). For *Halothece* sp., the PS beads did not cause aggregation of the cells, but a clear direct interaction was detected, with cyanobacteria starting to engulf the beads (Figs. 5F and 5G). For *Cobetia* sp., cells and the PS beads formed large aggregates (Figs. 5H and 5I).

**Figure 5.**
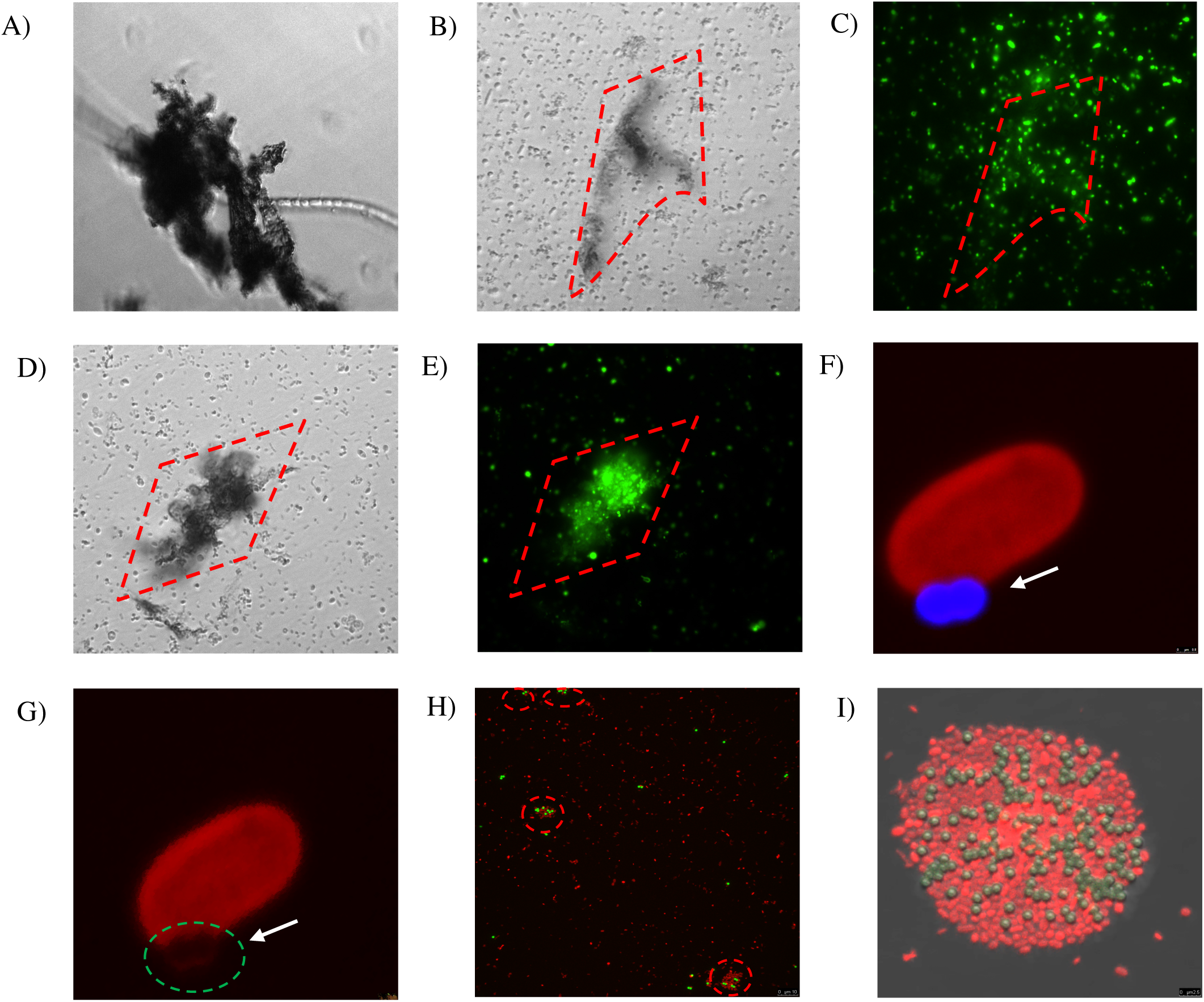
Direct physical interaction between MPs and bacterial species tested in which **A)** to **E)** show the interaction between diazotrophs and mix of MPs (PE, PP and PVC) at 100 μg mL^-1^ through inverted microscopy: **A)** *Fischerella muscicola* under bright-field (BF), **B)** and **C)** *Cobetia* sp. under BF and Sybr-green channel, respectively, **D)** and **E)** *Marinobacterium litorale* under BF and Sybr-green channel, respectively; **F)** to **I**) show the physical interaction between diazotrophs and PS 1 μm beads (excitation of 441 nm and emission of 485 nm) through confocal microscopy: **F)** and **G)** *Halothece* sp. under excitation of 532 nm wavelength and emission of 555-619 nm. The arrows and green dashed circle show where the cell membrane is interacting with the PS bead (blue). **H)** and **I)** *Cobetia* sp. under excitation of 493 nm and 636 nm of emission. The dashed red circles show cell agglomeration on the PS beads (yellow). Images were taken at 1000x (**A**, **B**, **C**, **D**, **E** and **H**) and further magnified (**F**, **G** and **I**).

### 3.3 Effect of varying concentrations of MPs and plastic additives in the phosphorus acquisition mechanisms of the diazotrophs

The MPs and their plastic additives generally enhanced the alkaline phosphatase activity (APA) of *Halothece* sp., particularly with the addition of PS beads (at 4.55 x 10^7^ particles mL^-1^), reaching a maximum rate of reaction (V^max^) of 0.21 fmol MUF cell^-1^ h^-1^, significantly higher than controls (V^max^ = 0.033 fmol MUF cell^-1^ h^-1^) (Bonferroni test, p < 0.05, Fig. 6A). For *Halothece* sp., PO_4_^3-^-uptake rates were significantly downregulated by the addition of MPs and their plastic additives (ANOVA or Kruskal-Wallis, p < 0.05, Fig. 6B).

**Figure 6.**
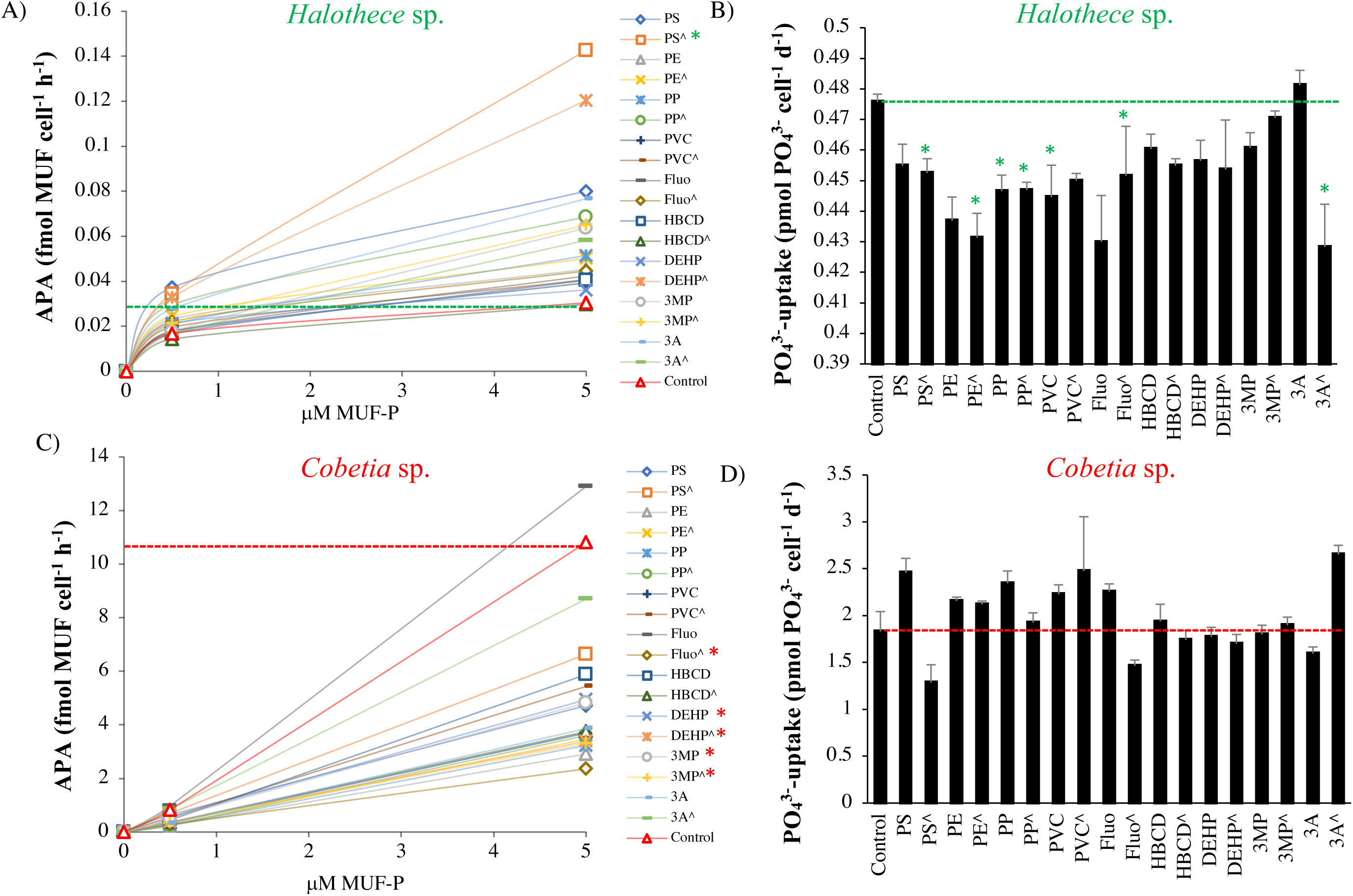
Phosphorus acquisition mechanisms for two species of N_2_-fixers exposed to different MPs and their plastic additives: **A)** Alkaline phosphatase activity (APA) and **B)** PO_4_^3-^-uptake for *Halothece* sp. **C)** APA and **D)** PO_4_^3-^-uptake for *Cobetia* sp. (^) indicate treatments up to 1000 μg mL^-1^ (MPs) and 3000 μg L^-1^ (organic additives), and without (^) indicate treatments 100 μg mL^-1^ (MPs) and 300 μg L^-1^ (organic additives). Values are the mean and the error bar is the standard error between the replicates. Asterisks (*) indicate significant differences (p < 0.05) compared with the controls, using a posthoc test (Bonferroni or Dunn test) after ANOVA or Kruskal-Wallis over the whole dataset.

Unlike *Halothece* sp., APA for *Cobetia* sp. was generally reduced by MPs and their organic additives (ANOVA, p < 0.05, Fig. 6C). Among the MPs, PE addition at high concentrations caused the most significant decrease in APA (V^max^ = 8.52 fmol MUF cell^-1^ h^-1^) compared to controls (V^max^ = 32.78 fmol MzzzUF cell^-1^ h^-1^) (Bonferroni test, p < 0.05, Fig. 6C). Among the plastic additives, fluoranthene caused the highest decrease in APA (V^max^ = 11.52 fmol MUF cell^-1^ h^-1^) compared to controls (Bonferroni test, p < 0.05, Fig. 6C). However, no significant differences in PO_4_^3-^-uptake rates were observed among all the treatments tested (ANOVA or Kruskal-Wallis, p > 0.05, Fig. 6D).

### 3.4 Effect of varying concentrations of MPs and plastic additives in N_2_-fixation

Specific N_2_-fixation rates (nmol N_2_-fixed cell^-1^ h^-1^) for *Halothece* sp. did not vary significantly with the addition of MPs nor their plastic additives (ANOVA or Kruskal-Wallis, p > 0.05, Fig. 7).

**Figure 7.**
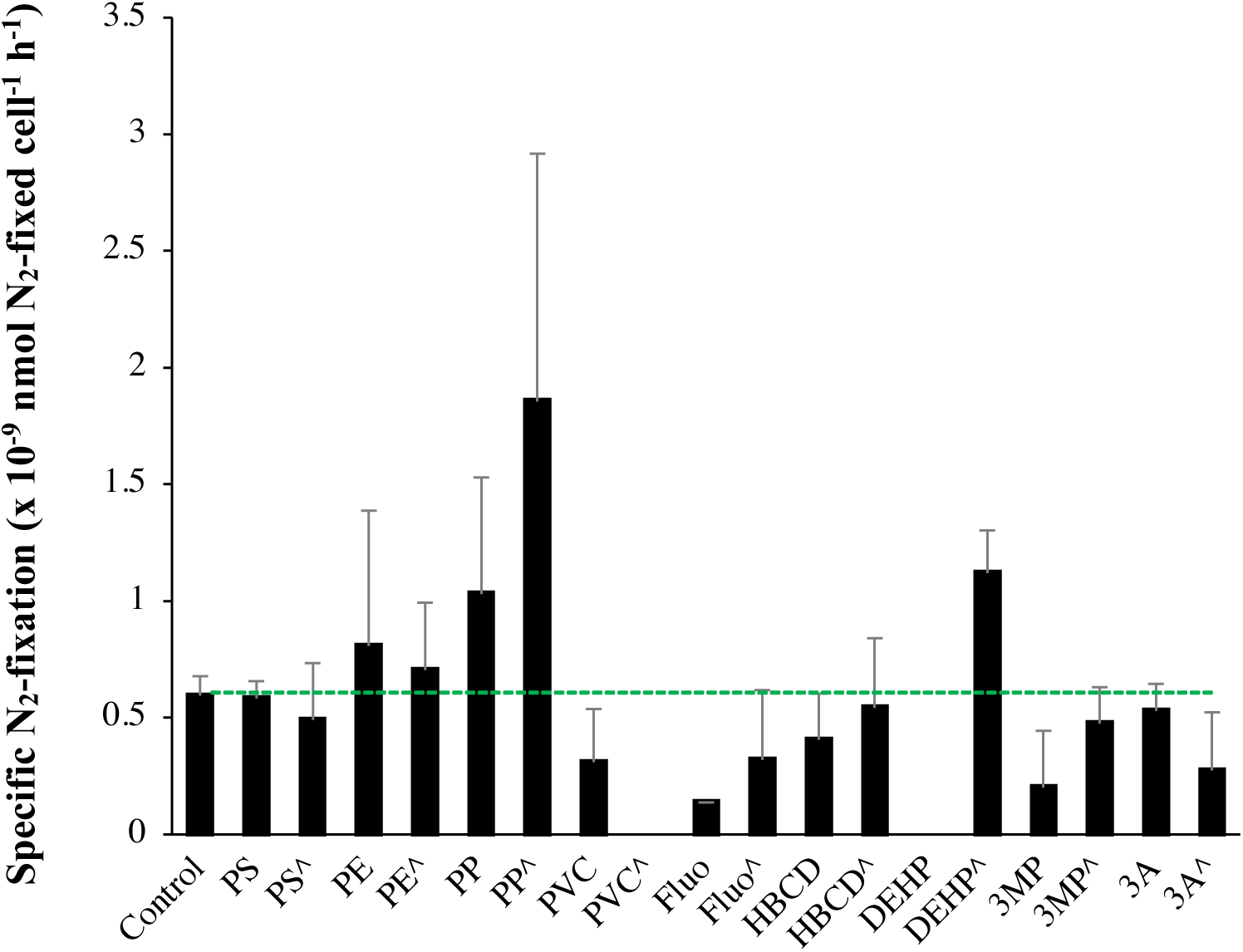
Specific N_2_-fixation rates (nmol N_2_-fixed cell^-1^ h^-1^) in the unicellular cyanobacteria *Halothece* sp. exposed to MPs and their plastic additives. (^) indicate treatments up to 1000 μg mL^-1^ (MPs) and 3000 μg L^-1^ (organic additives), and without (^) indicate treatments with 100 μg mL^-1^ (MPs) and 300 μg L^-1^ (organic additives). Values are the mean and the error bar is the standard error between the replicates.

## 4. Discussion

This is the first study under *in vitro* controlled conditions, in which the effects of the most predominant MPs (*e.g.* PE, PP, PVC and PS) in the oceans and their commonly associated organic additives (*i.e.* fluoranthene, HBCD and DEHP) were investigated in marine N_2_-fixing bacteria. Our results show that the responses were dependent on the concentrations of MPs and additives and that different species of N_2_-fixing bacteria with different sizes also varied in their response depending on the type of MPs, organic additives and their interacting effects. These effects can be beneficial (the “good”), deleterious (the “bad”) or both (the “double-sword”) (Fig. 8).

**Figure 8.**
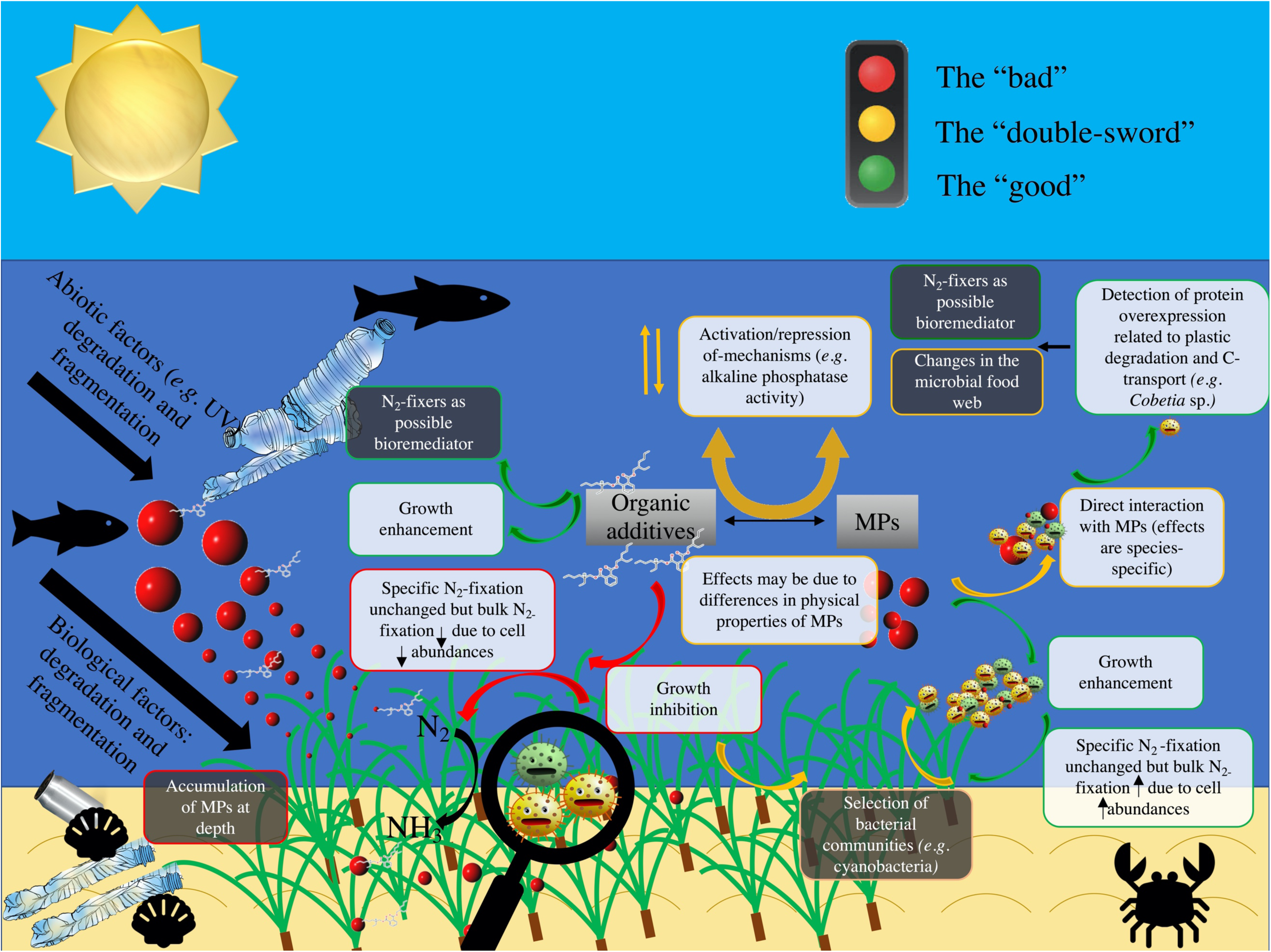
Effects of exposure of MPs and their organic additives on diazotrophs associated with seagrasses (e.g. *Posidonia oceanica*). The effects can be beneficial or “the good” (green arrows), deleterious or “the bad” (red arrows) and “double-sword” (yellow arrows). White boxes represent the results obtained in the present study, while the black boxes represent the implications of our results. MPs and their organic additives can trigger species-specific responses, and these responses are suggested to be dependent on the physical properties of the different MPs tested. The species that were able to grow with particular types of MP/additives can be possible bioremediators and the results also imply the selection for specific bacterial communities and changes in the microbial food web.

### 4.1 Effect of increasing concentrations of MPs and organic additives on N_2_-fixers

The importance of the concentration of MPs and organic additives in inducing significant effects on N_2_-fixers has been shown here when our test species, *e.g. Halothece* sp. and *Cobetia* sp. generally responded significantly to higher concentrations of MPs (Figs. 3A-C) compared with lower concentrations (Figs. 1A-F and 2). The importance of dose-dependent exposure to pollutants in affecting marine microbial populations is also exemplified in a pioneering study by Tetu et al. (2019) which suggests that increasing levels of leachates from PP and PVC (with unknown additives) can affect negatively the most abundant prokaryote in the marine environment, *Prochlorococcus*. This suggests the need to determine the thresholds levels of MPs and additives concentrations starting from which significant effects can be observed for key microbial populations in marine systems, and these data are necessary for effective environmental quality control management.

The ranges of MPs and organic additives concentrations added in the first part of our study (0 to 100 μg mL^-1^ and 0 to 300 μg L^-1^, respectively) were environmentally relevant and simulating observable levels in the marine environment. Although available data from the Mediterranean Sea reported low MPs concentrations of surface water (0 to 0.026μg mL^-1^, Suaria et al. 2016), sediment concentrations can be potentially higher. Indeed, a recent study points that MPs in the seafloor of the oceans can accumulate up to 191 pieces of MPs per 50 g of dried sediment (Kane et al., 2020), and considering the average weight of MPs can be 0.005 mg (Claessens et al., 2011) and the density of dry sediment is around 1500 Kg m^-3^, the seafloor can accumulate up to 29 μg mL^-1^ of MPs. Other marine environments, *e.g.* beaches, can potentially accumulate up to 60.38 μg mL^-1^ (1580 to 8050 particles Kg^-1^ dry sediment) (Everaert et al., 2018) and even up to 133 μg mL^-1^ (89 mg particles Kg^-1^ dry sediment) in hot-spots areas (Reddy et al., 2006). The concentrations of organic additives of MPs that can be found in seawater can accumulate up to 23.42 μg L^-1^ (*i.e.* phthalates) and in dry sediment up to 149 μg L^-1^ (Hermabessiere et al., 2017). In extremely impacted areas, organic additives of MPs can reach up to 2988 μg L^-1^ (Hermabessiere et al., 2017). Thus, in the second part of the experiment wherein we added up to 1000 μg mL^-1^ of MPs and up to 3000 μg L^-1^ of organic additives, we simulated “the worst-case scenario” considering extremely impacted areas (*e.g.* great Pacific Garbage Patch) (Lebreton et al., 2018).

### 4.2 Species-specific responses of N_2_-fixers with the addition of MPs and their implications

Species-specific growth responses of N_2_-fixers with the addition of MPs were exemplified here when PVC addition (at 100 µg mL^-1^) significantly enhanced the growth of autotrophic cyanobacterial diazotrophs (*Halothece* sp. and *F. muscicola*) and not on the three heterotrophic diazotrophs (*Cobetia* sp., *M. litorale* and *P. azotifigens*) (Fig. 1C). The enhancement of the growth of cyanobacterial diazotrophs (and the heterotrophic strain, *Cobetia* sp., only under high concentration, Figs. 1A-C and 3A) after 72 h of incubation can be due to increased dissolved organic carbon (DOC) pool in the medium that could have leached from the MPs added. This suggestion follows from Romera-Castillo et al. (2018) who reported that instantaneous leaching of DOC occurs right after the addition of MPs to seawater and 60% of the leached DOC can be bioavailable in < 120 h and can be rapidly consumed by microbes. The positive response of cyanobacterial diazotrophs (*Halothece* sp. and *F. muscicola*) with PVC addition also implies that these species may also be mixotrophs (DOC uptake occurs simultaneously with photosynthesis) and have broad implications for competition for DOC with the heterotrophs and organic carbon fluxes within the microbial food web. Still with unknown reasons, the cyanobacterial N_2_-fixers may have faster DOC uptake rates than the heterotrophs. It may take very high concentrations of leached DOC from high concentrations of MPs for heterotrophic diazotrophs to significantly enhance their growth rate as suggested by the increased growth of heterotrophic *Cobetia* sp. after exposure to high concentrations of MPs (Fig. 3A) and not in lower concentrations (Figs. 1A-C).

Species-specific responses based on protein expression profile are also shown here when two proteins related to plastic degradation (alcohol dehydrogenase [ADH]) and carbon transport (C4-dicarboxylate ABC transporter substrate-binding protein [C4-ABCS]) were overexpressed in the heterotroph *Cobetia* sp. and not in the cyanobacteria *Halothece* sp. after the addition of MPs at high concentrations (Fig. 3A). ADHs from *Rhodopseudomonas acidophila* M402 and *Pseudomonas oleovorans* have been shown to oxidize plastic polymers (*e.g.* PEG), being NAD dependent (Kawai, 2010; Ohta et al., 2006), consistent with our *in silico* structural analysis for *Cobetia* sp. (Figs. 4A-C). Further experimental studies have to be performed to evaluate if *Cobetia* sp. can actually degrade PEG polymers or similar ones, and indeed if the carbon released by ADH is transported by C4-ABCS inside the cells. Based on our initial results of protein expression profiles, some species of marine N_2_-fixers, aside from their important ecological role of providing new N into marine ecosystems, may have another important environmental role of biodegrading synthetic plastic polymers into smaller molecules, using it as a C-source for growth.

### 4.3 Size-specific responses of N_2_-fixers and MPs substrate-specific effects

Aside from the species-specific responses of N_2_-fixers after addition of MPs, we also report on the importance of size of the MPs and the size of the bacterial cells themselves in modulating the response of the diazotrophs. Larger-sized MPs enhanced the growth of autotropic and heterotrophic N_2_-fixers at high concentrations of MPs (Fig. 3A). Larger-sized MPs may provide more surface area for the cells to adhere as observed in the heterotrophs tested here (Figs. 5B-E), and surface attachment of the cells may facilitate nutrient uptake, through which necessary metabolites and co-factors can be obtained as suggested by Tuson and Weibel (2013). Smaller MPs (*i.e.* PS beads of 1 µm-size), however, affected negatively the smallest size heterotrophic bacteria (*i.e. Cobetia* sp. of~ 1 µm-size), and not the unicellular cyanobacteria (*i.e. Halothece* sp. of ~ 4 to 7 μm-size) (Fig. 3C). This may be due to the differences in the degree of physical adhesion between the PS beads and different species of N_2_-fixers. Approximately 40 to 87% of *Cobetia* sp. cells were adhered to PS (Figs. 3C and 5H-I), while only ~ 2 to 5% of *Halothece* sp. cells were adhered (Fig. 3C). Although less PS beads adhered on *Halothece* sp. cells, an invagination of the cell membranes by PS beads has been observed (Figs. 5F and 5G). The mechanisms of these observed physical adhesion and their effect on cell growth need to be further investigated.

The physicochemical properties (*e.g.* hydrophobicity, electrostatic attraction or roughness) of different MPs may affect the responses of bacterial communities (Ogonowski et al., 2018). Hydrophobicity, for example, can directly affect the bacterial colonization on MPs. PS polymers which have aromatic phenyl groups are one of the most hydrophobic polymers (Ogonowski et al., 2018), and this may explain the adherence of both autotrophic and heterotrophic N_2_-fixers to PS beads (Figs. 5F-I). However, only the heterotrophic N_2_-fixers (*Cobetia* sp., *M. litorale* and *P. azotifigens*) tested were able to adhere on other types of MPs (*i.e.* PE, PP and PVC, Figs. 5A-E) which are less hydrophobic than PS. Moreover, PVC polymers are slightly more hydrophilic than PE (Kennedy, 2014), suggesting that they can be less available for adhesion and more available for bacterial growth. This may explain why PVC polymers were the MPs that most enhanced bacteria growth (Figs. 1C, 2 and 3A). Further studies are needed to investigate if indeed hydrophobicity of MPs can affect the growth responses of bacterial cells.

### 4.4 Effect of organic additives on growth of N^2^-fixers

The effect of organic additives on the growth of N_2_-fixers is suggested here to be dependent on the type of additive. The additive DEHP for example, had a positive effect on the growth of heterotrophic *Cobetia* sp. indicating that this species might use DEHP as a C-source (Fig. 3B) and can be a possible bioremediator of DEHP-contaminated environments like *Rhodococcus* (Wang et al., 2015). The polycyclic aromatic hydrocarbon (PAH) additive, fluoranthene, on the other hand, had a negative effect on heterotrophic *Cobetia* sp. and on the autotrophs (*Halothece* sp. and *F. muscicola*, Figs. 1D and 3B), consistent with previous toxicity studies done on larger microorganisms (*e.g.* the marine diatom *Phaeodactylum tricornutum,* Wang and Zheng, 2008). The differing sensitivities of different species of N_2_-fixers to a particular type of additive can be due to cell-size dependent toxicity. For example, here we found an increasing toxicity to a PAH additive, fluoranthene, from bigger-to smaller-sized N_2_-fixers: *Cobetia* sp. (~ 1 μm) > unicellular cyanobacteria *Halothece* sp. (4-7 μm), > filamentous cyanobacteria *F. muscicola* (~ 7 μm) (Figs. 1D and 3B). The negative correlation between cell size and PAH toxicity is consistent with the study of Echeveste et al. (2010), and may be due to higher surface to volume ratio of smaller-sized cells which increases the potential of the additives to be adsorbed or consumed by the cells. The mechanisms behind the responses of N_2_-fixers to different types of additives need to be further studied, particularly for fluoranthene which is reported to be degraded by bacteria isolated from the marine environment (Cao et al., 2015).

The responses of N_2_-fixers to the addition of MPs with their organic additives (*i.e.* beneficial effect, PVC + DEHP or detrimental, PE + fluoranthene) were more evident than the addition of MPs or additives alone (Fig. 2). This suggests a synergistic or antagonistic interaction of the pollutants. In the case of PVC + DEHP, their positive effect for the heterotrophs *Cobetia* sp. and *P. azotifigens* can be due to (I) the affinity of organic additives to MPs (MPs can sorb additives), making these chemical compounds less available for the cells (and thus less harmful) (Hahladakis et al., 2018), and (II) additives (*e.g.* DEHP, Fig. 3B) can be used as a C-source together with MPs (*e.g.* PVC, Fig. 3A), in a synergistic way. However, additives can be sorbed and/or liberated by MPs in the marine environments (Gallo et al., 2018, Hahladakis et al., 2018), increasing their availability to the bacteria and consequently affecting bacterial growth negatively, consistent with our results with PE + fluoranthene (Fig. 2).

### 4.5 Effect of MPs and organic additives on the P-mechanisms of N_2_-fixers

Generally, alkaline phosphatase activity (APA) is stimulated when dissolved inorganic phosphorus (DIP) concentrations are limiting, releasing inorganic phosphorus from dissolved organic phosphorus (DOP) (Santos-Beneit, 2015). In previous experiments, we described that cyanobacterial N_2_-fixer *Halothece* sp. synthesizes an alkaline phosphatase D (PhoD) that is Fe dependent (Fe as metal co-factor) (Fernández-Juárez et al., 2019). Since metals (*i.e*. Fe) can be accumulated onto the plastics (Rochman et al., 2014), it is hypothesized that MPs may promote an environment rich in Fe co-factors. Considering the P-dependence of important processes (*e.g.* N_2_-fixation, Fernández-Juárez et al., 2019), stimulation of APA can promote the growth of *Halothece* sp. The decreased PO_4_^3-^-uptake rates observed in *Halothece* sp. (Fig. 6B) may be due to the adsorption of phosphate ions by PE and PVC (Hassenteufel, 1963). Contrary to *Halothece* sp., the heterotrophic *Cobetia* sp. showed a decrease of its APA rates with additions of MPs and their additives (Fig. 6C) and resulted in no significant effects on PO_4_^3-^-uptake rates (Fig. 6D). This suggests species-specific differences of P-mechanisms and P-requirements and the responses of these processes to MPs and their additives. Further studies are needed to investigate the molecular processes behind these responses.

### 4.6 Effect of MPs and their additives on N_2_-fixation rates

In a seminal paper, Bryant et al. (2016) claimed that MPs may be hot spots of N_2_-fixing autotrophic bacteria, based on the high abundances of N_2_-fixation genes (*nifH*, *nifD* and *nifK*) in the metagenomes associated with the plastic. Unfortunately, the authors did not measure the N_2_-fixation activities. With this pioneering study, the effects of MPs and their additives on N_2_-fixation rates are reported for the first time in cyanobacteria (*i.e. Halothece* sp., Fig. 7). Our results reveal that MPs and their additives did not have a significant effect on specific N_2_-fixation rates of *Halothece* sp. However, growth was positively enhanced with the addition of MPs (*i.e.* PP, PE, PVC and PS, Fig. 3A), increasing cell abundances of the N_2_-fixers and thus indirectly enhancing bulk N_2_-fixation in the medium. The organic additives which showed to be detrimental to growth of *Halothece* sp. (*i.e.* fluoranthene, HBCD and DEHP, Figs. 1D and 3B) could have an opposite effect.

Further studies are needed to evaluate the potential impacts of increasing levels of MPs on N_2_-fixation rates as rates can be affected in three ways: (I) P-scavenging of MPs can inhibit N_2_-fixation since N_2_-fixation is P-dependent (Fernández-Juárez et al., 2019); (II) Degradation of MPs by certain species may lower their N_2_-fixation rates since changes in energy allocation of the cells may occur toward degradation of MPs than for the energy-costly process of N_2_-fixation; (III) Production of radical oxygen species (ROS) in N_2_-fixers triggered by MPs and their additives (López-Alforja et al., in prep.) can inhibit N_2_-fixation rates since the nitrogenase complex is sensitive to ROS and redox processes (Alquéres et al., 2010).

### 4.7 Environmental implications of the effects of MPs on N_2_-fixers found in association with Posidonia oceanica

Some bacterial communities can be naturally found attached to MPs, in the so-called plastisphere (Erni-Cassola et al., 2019; Kirstein et al., 2019; Zettler et al., 2013), and Bryant et al. (2016) reported a core group of microbes associated with small plastic fragments on oligotrophic surface waters (*e.g.* in the Mediterranean Sea). The community composition of the plastic-associated microbes can differ from that of their water column-counterparts (Bryant et al., 2016; Oberbeckmann et al., 2014; Zettler et al., 2013), supporting the idea that MPs may be selecting bacterial communities in the ocean. Based on our results, changes in the composition of the microbiota (*e.g.* N_2_-fixing bacteria) may be occurring in *P. oceanica* meadows since plastic debris can accumulate in these meadows (Erni-Cassola et al., 2019; Reisser et al., 2015). Considering the beneficial effect of MPs on diazotrophic cyanobacterial growth more than on the heterotrophs, the selection for the dominance of the diazotrophic cyanobacteria may take place in *P. oceanica*. The consequences for this possible shift of diazotrophic community composition on *P. oceanica* should be further studied. In contrast with the effects of MPs, the organic additives, with the exception to those that can be degraded, can have detrimental effects on the diazotrophs and consequently affecting the maintenance of the productivity of *P. oceanica* since the activity of these N_2_-fixers can provide up to 100% of the N demand of the plants (Agawin et al., 2016). In addition, different physical processes such as currents or eddies may transport both MPs and the diazotrophs in adjacent seawater environments, since these cells can adhere with MPs (Figs. 5B-I), allowing them to settle and colonize other ecological niches. The role of MPs as vectors for transport of microorganisms has already been reported for pathogenic bacteria and harmful algal bloom-forming species (HABs) (Curren and Leong, 2019; Naik et al., 2019).

In summary, this study represents the first one to be conducted to investigate the direct effects of exposure to MPs and organic additives in N_2_-fixing bacteria found in association with seagrasses. We reported species-specific responses to MPs (*i.e.* PE, PP, PVC and PS) and organic additives (*i.e.* fluoranthene, HBCD and DEHP). Further studies should consider investigating in detail the mechanisms and processes behind these responses. The use of next generation analysis (*i.e.* transcriptomic or proteomic assays) to properly identify changes in gene expression or protein profiles derived from MPs and plastic additives will be helpful to allow a better comprehension of the molecular responses behind the plastic threat in the oceans.

## 6. Author Contributions

VFJ and XLA conducted all experiments with the help of AFC, PE, ABF, GRM, RMG in the various parameters measured in the study. NSRA is the supervisor of the laboratory.

## 8. Acknowledgments

We acknowledge the support and help of scientific technical service (Maria Trinidad Garcia Barceló) of the University of the Balearic Islands for gas-chromatography analyses. We also thank Pere Ferriol Buñola and Alba Coma Ninot for the help in the acquisition of the cultures, and project AAEE117/2017 of the Dirección General Innovación y Recerca CAIB.

## References

1. Agawin, N.S.R., Ferriol, P., Sintes, E., 2019. Simultaneous measurements of nitrogen fixation activities associated with different plant tissues of the seagrass *Posidonia oceanica*. Mar. Ecol. Prog. Ser. https://doi.org/10.3354/meps12854

2. Agawin, N.S.R., Benavides, M., Busquets, A., Ferriol, P., Stal, L.J., Arístegui, J., 2014. Dominance of unicellular cyanobacteria in the diazotrophic community in the Atlantic Ocean. Limnol. Oceanogr. 59, 623–637. https://doi.org/10.4319/lo.2014.59.2.0623

3. Agawin, N.S.R., Ferriol, P., Cryer, C., Alcon, E., Busquets, A., Sintes, E., Vidal, C., Moyà, G., 2016. Significant nitrogen fixation activity associated with the phyllosphere of Mediterranean seagrass *Posidonia oceanica*: First report. Mar. Ecol. Prog. Ser. 551, 53–62. https://doi.org/10.3354/meps11755

4. Agawin, N.S.R., Ferriol, P., Sintes, E., Moyà, G., 2017. Temporal and spatial variability of in situ nitrogen fixation activities associated with the Mediterranean seagrass *Posidonia oceanica* meadows. Limnol. Oceanogr. 62, 2575–2592. https://doi.org/10.1002/lno.10591

5. Almroth, B.C., Eggert, H., 2019. Marine plastic pollution: sources, impacts, and policy issues, Review of Environmental Economics and Policy, 13, 2, 317–326, https://doi.org/10.1093/reep/rez012

6. Alquéres, S.M.C., Oliveira, J.H.M., Nogueira, E.M., Guedes, H. V., Oliveira, P.L., Câmara, F., Baldani, J.I., Martins, O.B., 2010. Antioxidant pathways are up-regulated during biological nitrogen fixation to prevent ROS-induced nitrogenase inhibition in *Gluconacetobacter diazotrophicus*. Arch. Microbiol. 192, 835–841. https://doi.org/10.1007/s00203-010-0609-1

7. Anderson, O.R., 2018. Evidence for coupling of the carbon and phosphorus biogeochemical cycles in freshwater microbial communities. Front. Mar. Sci. 5, 1– 6. https://doi.org/10.3389/fmars.2018.00020

8. Browning, T.J., Achterberg, E.P., Yong, J.C., Rapp, I., Utermann, C., Engel, A., Moore, C.M., 2017. Iron limitation of microbial phosphorus acquisition in the tropical North Atlantic. Nat. Commun. 8, 1–7. https://doi.org/10.1038/ncomms15465

9. Bryant, J.A., Clemente, T.M., Viviani, D.A., Fong, A.A., Thomas, K.A., Kemp, P., Karl, D.M., White, A.E., DeLong, E.F., 2016. Diversity and activity of communities inhabiting plastic debris in the North Pacific Gyre. mSystems, 1, 1– 19. https://doi.org/10.1128/msystems.00024-16

10. Cao, J., Lai, Q., Yuan, J., Shao, Z., 2015. Genomic and metabolic analysis of fluoranthene degradation pathway in *Celeribacter indicus* P73 T. Sci. Rep. 5, 1– 12. https://doi.org/10.1038/srep07741

11. Campagne, C. S., Salles, J. M., Boissery, P., and Deter, J., 2015. The seagrass *Posidonia oceanica*: ecosystem services identification and economic evaluation of goods and benefits. Mar. Pollut. Bull. 97, 391–400. doi: 10.1016/j.marpolbul.2015.05.061

12. Claessens, M., Meester, S. De Landuyt, L. Van Clerck, K. De Janssen, C.R., 2011. Occurrence and distribution of microplastics in marine sediments along the Belgian coast. Mar. Pollut. Bull. 62, 2199–2204. https://doi.org/10.1016/j.marpolbul.2011.06.030

13. Cole, M., Galloway, T.S., 2015. Ingestion of nanoplastics and microplastics by pacific oyster larvae. Environ. Sci. Technol. 49, 14625–14632. https://doi.org/10.1021/acs.est.5b04099

14. Curren, E., Leong, S.C.Y., 2019. Profiles of bacterial assemblages from microplastics of tropical coastal environments. Sci. Total Environ. 655, 313–320. https://doi.org/10.1016/j.scitotenv.2018.11.250

15. de Haan, W.P., Sanchez-Vidal, A., Canals, M., 2019. Floating microplastics and aggregate formation in the Western Mediterranean Sea. Mar. Pollut. Bull. 140, 523–535. https://doi.org/10.1016/j.marpolbul.2019.01.053

16. DeLano, W.L., 2002. PyMOL: An open-source molecular graphics tool. CCP4 Newsl. Protein Crystallogr. https://doi.org/10.1038/s41598-017-03842-2

17. Echeveste, P., Agustí, S., Dachs, J., 2010. Cell size dependent toxicity thresholds of polycyclic aromatic hydrocarbons to natural and cultured phytoplankton populations. Environ. Pollut. 158, 299–307. https://doi.org/10.1016/j.envpol.2009.07.006

18. Erni-Cassola, G., Wright, R.J., Gibson, M.I., Christie-Oleza, J.A., 2019. Early colonization of weathered polyethylene by distinct bacteria in marine coastal seawater. Microb. Ecol. https://doi.org/10.1007/s00248-019-01424-5

19. Erni-Cassola, G., Zadjelovic, V., Gibson, M.I., Christie-oleza, J.A., 2019. Distribution of plastic polymer types in the marine environment ; A meta-analysis. J. Hazard. Mater. 369, 691–698. https://doi.org/10.1016/j.jhazmat.2019.02.067

20. Everaert, G., Van Cauwenberghe, L., De Rijcke, M., Koelmans, A.A., Mees, J., Vandegehuchte, M., Janssen, C.R., 2018. Risk assessment of microplastics in the ocean: Modelling approach and first conclusions. Environ. Pollut. 242, 1930–1938. https://doi.org/10.1016/j.envpol.2018.07.069

21. Fernández-Juárez, V., Bennasar-Figueras, A., Tovar-Sanchez, A., Agawin, N.S.R, 2019. The role of iron in the P-acquisition mechanisms of the unicellular N_2_-fixing cyanobacteria *Halothece* sp ., found in association with the Mediterranean seagrass *Posidonia oceanica*. Front. Microbiol. 10, 1–22. https://doi.org/10.3389/fmicb.2019.01903

22. Finn, R.D., Coggill, P., Eberhardt, R.Y., Eddy, S.R., Mistry, J., Mitchell, A.L., Potter, S.C., Punta, M., Qureshi, M., Sangrador-Vegas, A., Salazar, G.A., Tate, J., Bateman, A., 2016. The Pfam protein families database: Towards a more sustainable future. Nucleic Acids Res. 44, D279–D285. https://doi.org/10.1093/nar/gkv1344

23. Gallo, F., Fossi, C., Weber, R., Santillo, D., Sousa, J., Ingram, I., Nadal, A., Romano, D., 2018. Marine litter plastics and microplastics and their toxic chemicals components: the need for urgent preventive measures. Environ. Sci. Eur. 30. https://doi.org/10.1186/s12302-018-0139-z

24. Ghaffar, S., Stevenson, R.J., Khan, Z., 2017. Effect of phosphorus stress on *Microcystis aeruginosa* growth and phosphorus uptake. PLoS One, 12. https://doi.org/10.1371/journal.pone.0174349

25. Grosdidier, A., Zoete, V., Michielin, O., 2011. SwissDock, a protein-small molecule docking web service based on EADock DSS. Nucleic Acids Res. 39, 270–277. https://doi.org/10.1093/nar/gkr366

26. Hahladakis, J.N., Velis, C.A., Weber, R., Iacovidou, E., Purnell, P., 2018. An overview of chemical additives present in plastics: Migration, release, fate and environmental impact during their use, disposal and recycling. J. Hazard. Mater. 344, 179–199. https://doi.org/10.1016/j.jhazmat.2017.10.014

27. Hansen, H.P, Koroleff, F., 1999. “Determination of nutrients” in method of seawater analysis. eds. K. Grasshoff, K. Kremling, and M. Ehrhardt. doi: 10.1002/9783527613984.ch10

28. Harrison P., Sapp., M., Schratzberger, M., Osborn, A.M., 2011. Interactions between microorganisms and marine microplastics: a call for re-search. Mar Technol Soc J. 45 (2), 12–20. https://doi.org/10.1093/jac/10.2.84

29. Hartmann, N.B., Hüffer, T., Thompson, R.C., Hassellöv, M., Verschoor, A., Daugaard, A.E., Rist, S., Karlsson, T., Brennholt, N., Cole, M., Herrling, M.P., Hess, M.C., Ivleva, N.P., Lusher, A.L., Wagner, M., 2019. Are we speaking the same language? Recommendations for a definition and categorization framework for plastic debris. Environ. Sci. Technol. 53, 1039–1047. https://doi.org/10.1021/acs.est.8b05297

30. Hassenteufel, W., Jagitsch, R., Koczy, F.F., 1963. Impregnation of glass surfacae against sortion of phospahate traces. Limnol. Oceanogr. 8, 152–156. https://doi.org/10.4319/lo.1963.8.2.0152

31. Hermabessiere, L., Dehaut, A., Paul-Pont, I., Lacroix, C., Jezequel, R., Soudant, P., Duflos, G., 2017. Occurrence and effects of plastic additives on marine environments and organisms: A review. Chemosphere, 182, 781-793 https://doi.org/10.1016/j.chemosphere.2017.05.096

32. Huang, B., 2009. MetaPocket: a meta approach to improve protein ligand binding site prediction. OMICS, 13 (4), 325-330. doi:10.1089/omi.2009.0045

33. Ivleva N.B., Golden, S.S., 2007. Protein extraction, fractionation, and purification from cyanobacteria. Methods in Molecular Biology, 362, 365–73. https://doi.org/10.1007/978-1-59745-257-1

34. Jaén-Luchoro, D., Aliaga-Lozano, F., Gomila, R.M., Gomila, M., Salvà-Serra, F., Lalucat, J., Bennasar-Figueras, A., 2017. First insights into a type II toxin-antitoxin system from the clinical isolate *Mycobacterium* sp. MHSD3, similar to epsilon/zeta systems. PLoS One, 12, 1–20. https://doi.org/10.1371/journal.pone.0189459

35. Jensen, B.B., Cox, R.P., 1983. Direct measurements of steady-state kinetics of cyanobacterial N_2_ uptake by membrane-leak mass spectrometry and comparisons between nitrogen fixation and acetylene reduction. Appl. Environ. Microbiol. 45, 1331–1337.

36. Kane, I.A., Clare, M.A., Miramontes, E., Wogelius, R., Rothwell, J.J., Garreau, P., Pohl, F., 2020. Seafloor microplastic hotspots controlled by deep-sea circulation. Science (80-.), 5899. doi: 1–11.10.1126/science.aba5899

37. Kawai, F., 2010. The biochemistry and molecular biology of xenobiotic polymer degradation by microorganisms. Biosci. Biotechnol. Biochem. 74 (9), 1743–1759. https://doi.org/10.1271/bbb.100394

38. Kennedy, E.J., 2014. Biological drug products: Development and strategies. Edited by Wei Wang and Manmohan Singh. ChemMedChem. https://doi.org/10.1002/cmdc.201402432

39. Kirstein, I.V., Wichels, A., Gullans, E., Krohne, G., Gerdts, G., 2019. The plastisphere – Uncovering tightly attached plastic “specific” microorganisms. PLoS One, 14, 1–17. https://doi.org/10.1371/journal.pone.0215859

40. Lebreton, L., Slat, B., Ferrari, F., Sainte-Rose, B., Aitken, J., Marthouse, R., Hajbane, S., Cunsolo, S., Schwarz, A., Levivier, A., Noble, K., Debeljak, P., Maral, H., Schoeneich-Argent, R., Brambini, R., Reisser, J., 2018. Evidence that the Great Pacific Garbage Patch is rapidly accumulating plastic. Sci. Rep. 8, 1–15. https://doi.org/10.1038/s41598-018-22939-w

41. Naik, R.K., Naik, M.M., D’Costa, P.M., Shaikh, F., 2019. Microplastics in ballast water as an emerging source and vector for harmful chemicals, antibiotics, metals, bacterial pathogens and HAB species: A potential risk to the marine environment and human health. Mar. Pollut. Bull. 149, 110525. https://doi.org/10.1016/j.marpolbul.2019.110525

42. Nelms, S.E., Galloway, T.S., Godley, B.J., Jarvis, D.S., Lindeque, P.K., 2018. Investigating microplastic trophic transfer in marine top predators. Environ. Pollut. 238, 999–1007. https://doi.org/10.1016/j.envpol.2018.02.016

43. Oberbeckmann, S., Loeder, M.G.J., Gerdts, G., Osborn, M.A., 2014. Spatial and seasonal variation in diversity and structure of microbial biofilms on marine plastics in Northern European waters. FEMS Microbiol. Ecol. 90, 478–492. https://doi.org/10.1111/1574-6941.12409

44. Ogonowski, M., Motiei, A., Ininbergs, K., Hell, E., Gerdes, Z., Udekwu, K.I., Bacsik, Z., Gorokhova, E., 2018. Evidence for selective bacterial community structuring on microplastics. Environ. Microbiol. 20, 2796–2808. https://doi.org/10.1111/1462-2920.14120

45. Ohta, T., Kawabata, T., Nishikawa, K., Tani, A., Kimbara, K., Kawai, F., 2006. Analysis of amino acid residues involved in catalysis of polyethylene glycol dehydrogenase from *Sphingopyxis terrae*, using three-dimensional molecular modeling-based kinetic characterization of mutants. Appl. Environ. Microbiol. 72, 4388–4396. https://doi.org/10.1128/AEM.02174-05

46. Paul-Pont, I., Tallec, K., Gonzalez-Fernandez, C., Lambert, C., Vincent, D., Mazurais, D., Zambonino-Infante, J.L., Brotons, G., Lagarde, F., Fabioux, C., Soudant, P., Huvet, A., 2018. Constraints and priorities for conducting experimental exposures of marine organisms to microplastics. Front. Mar. Sci. 5, 1–22. https://doi.org/10.3389/fmars.2018.00252

47. Reddy, M.S., Shaik Basha, Adimurthy, S., Ramachandraiah, G., 2006. Description of the small plastics fragments in marine sediments along the Alang-Sosiya ship-breaking yard, India. Estuar. Coast. Shelf Sci. 68 (3-4), 656–660. https://doi.org/10.1016/j.ecss.2006.03.018

48. Reisser, J., Slat, B., Noble, K., Plessis, K., Epp, M., Proietti, M., Sonneville, J. De Becker, T., 2015. The vertical distribution of buoyant plastics at sea: an observational study in the North Atlantic Gyre. Biogeosciences, 12, 1249–1256. https://doi.org/10.5194/bg-12-1249-2015

49. Renn, C.E., 1937. Bacteria and the phophorus cycle in the sea. Biol. Bull. 72, 190–195.

50. Roche, D.B., Buenavista, M.T., McGuffin, L.J., 2013. The FunFOLD2 server for the prediction of protein-ligand interactions. Nucleic Acids Res. 41, 303–307. https://doi.org/10.1093/nar/gkt498

51. Rochman, C.M., Hentschel, B.T., Teh, S.J., 2014. Long-term sorption of metals is similar among plastic types: Implications for plastic debris in aquatic environments. PLoS One, 9 (1) e85433. https://doi.org/10.1371/journal.pone.0085433

52. Romera-Castillo, C., Pinto, M., Langer, T.M., Álvarez-Salgado, X.A., Herndl, G.J., 2018. Dissolved organic carbon leaching from plastics stimulates microbial activity in the ocean. Nat. Commun. 9,1430. https://doi.org/10.1038/s41467-018-03798-5

53. Santos-Beneit, F., 2015. The Pho regulon: A huge regulatory network in bacteria. Front. Microbiol. 6. https://doi.org/10.3389/fmicb.2015.00402

54. Schwarzenbach, R.P., Gschwend, P.M., Imboden, D.M., 2002. Environmental Organic Chemistry. https://doi.org/10.1002/0471649643

55. Sohm, J.A., Webb, E.A., Capone, D.G., 2011. Emerging patterns of marine nitrogen fixation. Nat. Rev. Microbiol. 9, 499–508. https://doi.org/10.1038/nrmicro2594

56. Stal, L.J., 1988. Acetylene reduction technique for assay of nitrogenase. Methods Enzymol. 167, 474–484.

57. Stanier, R.Y., Deruelles, J., Rippka, R., Herdman, M., Waterbury, J.B., 1979. Generic assignments, strain histories and properties of pure cultures of cyanobacteria. Microbiology, 111, 1–61. https://doi.org/10.1099/00221287-111-1-1

58. Tetu, S.G., Sarker, I., Schrameyer, V., Pickford, R., Elbourne, L.D.H., Moore, L.R., Paulsen, I.T., 2019. Plastic leachates impair growth and oxygen production in *Prochlorococcus*, the ocean’s most abundant photosynthetic bacteria. Commun. Biol. 2, 184. https://doi.org/10.1038/s42003-019-0410-x

59. Tuson, H.H., Weibel, D.B., 2013. Bacteria-surface interactions. Soft Matter, 9, 4368– 4380. https://doi.org/10.1039/c3sm27705d

60. Urbanek, A.K., Rymowicz, W., Mirończuk, A.M., 2018. Degradation of plastics and plastic-degrading bacteria in cold marine habitats. Appl. Microbiol. Biotechnol. 102, 7669–7678. https://doi.org/10.1007/s00253-018-9195-y

61. Wang, J., Zhang, M.Y., Chen, T., Zhu, Y., Teng, Y., Luo, Y.M., Christie, P., 2015. Isolation and identification of a di-(2-ethylhexyl) phthalate-degrading bacterium and its role in the bioremediation of a contaminated soil. Pedosphere, 25, 202–211. https://doi.org/10.1016/S1002-0160(15)60005-4

62. Wang, L., Zheng, B., 2008. Toxic effects of fluoranthene and copper on marine diatom *Phaeodactylum tricornutum*. J. Environ. Sci. 20, 1363–1372. https://doi.org/10.1016/S1001-0742(08)62234-2

63. Yang, Y., Liu, G., Song, W., Ye, C., Lin, H., Li, Z., Liu, W., 2019. Plastics in the marine environment are reservoirs for antibiotic and metal resistance genes. Environ. Int. 123, 79–86. https://doi.org/10.1016/j.envint.2018.11.061

64. Zettler, E.R., Mincer, T.J., Amaral-Zettler, L.A., 2013. Life in the ‘plastisphere’: Microbial communities on plastic marine debris. Environ. Sci. Technol. 47, 7137– 7146. https://doi.org/10.1021/es401288x

65. Zhang, Y., 2008. I-TASSER server for protein 3D structure prediction. BMC Bioinformatics, 9, 40. https://doi.org/10.1186/1471-2105-9-40

