## Supplementary Table 1 and Figure 1 for "The “good, the bad and the double-sword” effects of exposure to MPs and their organic additives on N_2_-fixing bacteria"

**Supplementary Table 1.** List of all treatments performed in this study, divided in two parts: (**I**) Response to MPs at environmentally relevant concentrations and (**II**) Response to MPs at environmentally concentrations vs high concentrations ("the worst case scenario"). Abbreviations: PE, polyethylene; PP, polypropylene; PVC, polyvinyl chloride.; Fluo, fluoranthene; HBCD, 1,2,5,6,9,10-hexabromocyclododecane; DEHP, dioctyl-phthalate; PS, polystyrene. Response variables of each experiment are indicated on the table.

| Treatments: | **MPs (PE, PP and/or PVC)** | **Additive (A) (Fluoranthene, HBCD and/or DEHP)** | **MPs and A: PE+Fluo/PP+HBCD/ PVC+DEHP or all interacting** | **PS (beads)** |
|---|---|---|---|---|
| **I. Response to MPs at environmentally relevant concentrations** | | | | $4.55\ x10^{3}$ particles $mL^{-1}$<br><br>In *Halothece* sp. PCC 7418 (data did not show) |
| Response variable: growth and microscopic analysis | | | | |
| *Halothece* sp. PCC 7418 | 0, 0.01, 0.1, 1, 100 $\mu g\ mL^{-1}$ | 0, 0.3, 3, 30, 300 $\mu g\ L^{-1}$ | Low (L) MP (0.01 $\mu g\ ml^{-1}$) and Low (L) A (0.3 $\mu g\ L^{-1}$)<br><br>High (H) MP (100 $\mu g\ ml^{-1}$) and High (H) A (300 $\mu g\ L^{-1}$) | |
| *Fischerella muscicola* PCC 73103 | | | | |
| *Cobetia* sp. UIB 001 | | | | |
| *Marinobacterium litorale* DSM 23545 | | | | |
| *Pseudomonas azotifigens* DSM $17556^{T}$ | | | | |
| **II. Response to MPs at environmentally relevant concentrations vs high concentrations ("the worst case scenario")** | | | | |
| Response variable: growth, microscopic analysis, protein overexpression, alkaline phosphatase activity (APA), $PO_4^{3-}$-uptake and $N_2$-fixation | | | | |
| *Halothece* sp. PCC 7418 | 0, 100 and 1000 $\mu g\ mL^{-1}$ | 0, 300, 3000 $\mu g\ L^{-1}$ | 3MP (100 $\mu g\ mL^{-1}$) and 3A (300 $\mu g\ L^{-1}$) (all combined)<br><br>3MP (1000 $\mu g\ mL^{-1}$) and 3A (3000 $\mu g\ L^{-1}$) (all combined) | $4.55\ x10^{6}$ particles $mL^{-1}$<br><br>$4.55\ x10^{7}$ particles $mL^{-1}$ |
| *Cobetia* sp. UIB 001 | | | | |

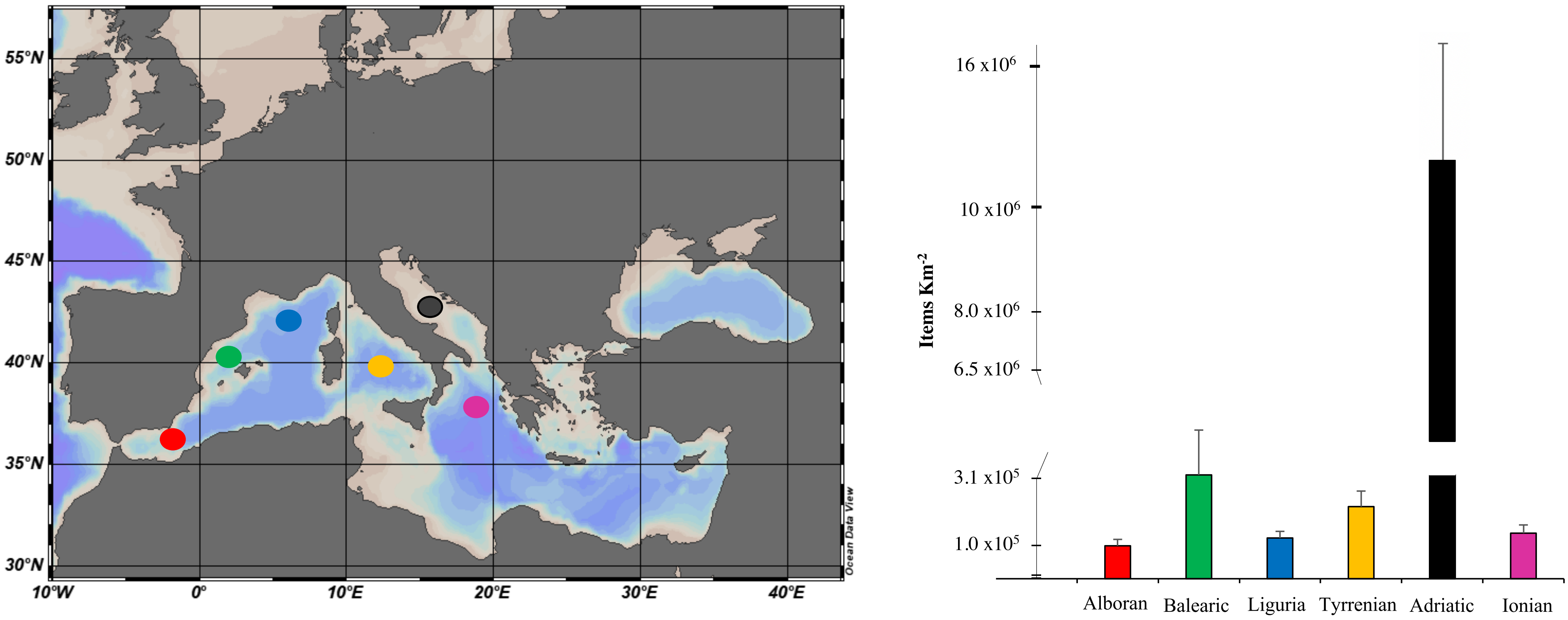
55°N
50°N
45°N
40°N
35°N
30°N
10°W
0°
10°E
20°E
30°E
40°E
Ocean Data View
Items Km$^{-2}$
16 x10$^{6}$
10 x10$^{6}$
8.0 x10$^{6}$
6.5 x10$^{6}$
3.1 x10$^{5}$
1.0 x10$^{5}$
Alboran
Balearic
Liguria
Tyrrenian
Adriatic
Ionian

**Supplementary Figure 1.** Marine plastic litter in the Mediterranean Sea. **A)** Items by $Km^{-2}$ are represented in the different seas of the Mediterranean Sea: Alboran Sea, Balearic Sea, Liguria Sea, Tyrrenian Sea, Adriatic Sea and Ionian Sea, represented by color. There were not data available for Aegean Sea. Data was extracted from https://litterbase.awi.de/litter from studies published between 2010 and 2019.

**Reference used to make supplementary Figure 1**


Collignon, A., Hecq, J. H., Glagani, F., Voisin, P., Collard, F., Goffart, A., 2012. Neustonic microplastic and zooplankton in the North Western Mediterranean Sea. Marine Pollution Bulletin, 64(4), 861–864. https://doi.org/10.1016/j.marpolbul.2012.01.011

de Haan, W. P., Sanchez-Vidal, A., Canals, M., 2019. Floating microplastics and aggregate formation in the Western Mediterranean Sea. Marine Pollution Bulletin, 140 (December 2018), 523–535. https://doi.org/10.1016/j.marpolbul.2019.01.053

Faure, F., Saini, C., Potter, G., Galgani, F., de Alencastro, L. F., Hagmann, P., 2015. An evaluation of surface micro- and mesoplastic pollution in pelagic ecosystems of the Western Mediterranean Sea. Environmental Science and Pollution Research, 22(16), 12190–12197. https://doi.org/10.1007/s11356-015-4453-3

Fossi, M. C., Romeo, T., Baini, M., Panti, C., Marsili, L., Campan, T., … Lapucci, C., 2017. Plastic debris occurrence, convergence areas and fin whales feeding ground in the Mediterranean marine protected area Pelagos Sanctuary: A modeling approach. Frontiers in Marine Science, 4(MAY), 1–15. https://doi.org/10.3389/fmars.2017.00167

Mistri, M., Infantini, V., Scoponi, M., Granata, T., Moruzzi, L., Massara, F., … Munari, C., 2017. Small plastic debris in sediments from the Central Adriatic Sea: Types, occurrence and distribution. Marine Pollution Bulletin, 124(1), 435–440. https://doi.org/10.1016/j.marpolbul.2017.07.063

Pedrotti, M. L., Petit, S., Elineau, A., Bruzaud, S., Crebassa, J. C., Dumontet, B., … Cózar, A., 2016. Changes in the floating plastic pollution of the Mediterranean Sea in relation to the distance to land. PLoS ONE, 11(8), 1–14. https://doi.org/10.1371/journal.pone.0161581

Ruiz-Orejón, L. F., Sardá, R., Ramis-Pujol, J., 2016. Floating plastic debris in the Central and Western Mediterranean Sea. Marine

Environmental Research, 120, 136–144. https://doi.org/10.1016/j.marenvres.2016.08.001

Ruiz-Orejón, L. F., Sardá, R., Ramis-Pujol, J., 2018. Now, you see me: High concentrations of floating plastic debris in the coastal waters of the Balearic Islands (Spain). Marine Pollution Bulletin, 133(January), 636–646. https://doi.org/10.1016/j.marpolbul.2018.06.010

Schmidt, N., Thibault, D., Galgani, F., Paluselli, A., Sempéré, R., 2018. Occurrence of microplastics in surface waters of the Gulf of Lion (NW Mediterranean Sea). Progress in Oceanography, 163, 214–220. https://doi.org/10.1016/j.pocean.2017.11.010

Vianello, A., Da Ros, L., Boldrin, A., Marceta, T., Moschino, V., 2018. First evaluation of floating microplastics in the Northwestern Adriatic Sea. Environmental Science and Pollution Research, 25(28), 28546–28561. https://doi.org/10.1007/s11356-018-2812-6

Zeri, C., Adamopoulou, A., Bojanić Varezić, D., Fortibuoni, T., Kovač Viršek, M., Kržan, A., … Vlachogianni, T., 2018. Floating plastics in Adriatic waters (Mediterranean Sea): From the macro- to the micro-scale. Marine Pollution Bulletin, 136 (July), 341–350. https://doi.org/10.1016/j.marpolbul.2018.09.016
